# Dynamic interplay between non-coding enhancer transcription and gene activity in development

**DOI:** 10.1101/2022.08.02.502573

**Authors:** Kota Hamamoto, Takashi Fukaya

## Abstract

Non-coding transcription at the intergenic regulatory regions is a prevalent feature of metazoan genomes, but its biological function remains uncertain. Here, we devised a live-imaging system that permits simultaneous visualization of gene activity along with intergenic non-coding transcription at the single-cell resolution in *Drosophila*. Quantitative image analysis revealed that elongation of RNA polymerase II across the internal core region of enhancers leads to suppression of transcriptional bursting from linked genes. Super-resolution imaging and genome-editing analysis further demonstrated that enhancer transcription antagonizes molecular crowding of transcription factors, thereby interrupting the formation of transcription hub at the gene locus. We also show that a certain class of developmental enhancers are structurally optimized to co-activate gene transcription together with non-coding transcription effectively. We suggest that enhancer function is flexibly tunable through the modulation of hub formation via surrounding non-coding transcription during development.

## Introduction

Enhancers are a class of regulatory DNAs that control spatial and temporal specificity of gene expression in development ^1^. Quantitative imaging and single-cell RNA transcriptome analyses revealed that enhancers mainly act to drive successive bursts of *de novo* RNA synthesis from their target genes ^2–4^. Recent whole-genome studies reported that non-coding transcription at the intergenic regulatory regions is a pervasive feature of metazoan genome ^5, 6^. Among these, non-coding enhancer transcription is thought to be a widespread mechanism conserved across species including mammals ^5–7^, flies ^8, 9^, and worms ^10^, and often used as a hallmark for identifying active enhancers in the genome ^e.g., 11^. A number of studies reported that the level of enhancer transcription correlates with the activity of nearby genes ^e.g., 5, 6, 12, 13^, implicating that that there is a functional interplay between these two reactions to facilitate gene expression. Whereas a variety of models have been proposed to explain molecular function of enhancer transcription so far ^reviewed^ ^in 14^, most of them stem from bulk and fixed analysis of heterogeneous population of cultured cells, and studies that directly link non-coding enhancer transcription and gene activity at the single-cell level are scarce. The fact that live visualization of non-coding enhancer transcription in multicellular organisms has been challenging hindered elucidation of how it can impact temporal dynamics of gene expression during animal development. In addition, it has also been implicated that non-coding transcription at the intergenic regulatory regions can lead to downregulation of nearby genes ^15–20^, which is seemingly incompatible with proposed positive regulatory functions of enhancer transcription. To reconcile these contradictory observations, there is a critical need for developing a new experimental framework that enables one to precisely modulate the mode of intergenic non-coding transcription and quantitatively visualize its functional impacts on gene activity in living multicellular organism. Here, we successfully devised a MS2/PP7 live-imaging system that permits simultaneous visualization of gene activity along with intergenic non-coding transcription in developing *Drosophila* embryos. Quantitative image analysis revealed that induction of transcriptional bursting from target genes is dramatically suppressed in the presence of enhancer transcription *in cis*. Using a variety of genome engineering and genetic approaches, we provide evidence that transcriptional attenuation takes place specifically when elongating RNA polymerase II (Pol II) transverses internal core region of enhancers. Super-resolution imaging and genome-editing analysis of key transcription factors Dorsal and Zelda further demonstrated that enhancer transcription counteracts molecular crowding of transcriptional activators to limit the formation of transcription hub at the gene locus. We also show that a certain class of developmental enhancers are structurally optimized to co-activate gene transcription together with non-coding transcription at the same time. We proposed that regulatory activities of developmental enhancers are flexibly tunable through the modulation of hub formation via surrounding non-coding transcription during development.

## Results

### Live-imaging of intergenic non-coding transcription in living embryos

In order to simultaneously monitor intergenic non-coding transcription along with gene activity at the single-cell resolution, we employed a newly developed MS2/PP7 two-color live-imaging method in *Drosophila* ^3, 21–23^. First, a sequence cassette containing 24x MS2 repeats was engineered into the 5’ untranslated region of the *yellow* reporter gene (Figure 1A; top). A well-characterized 1.5-kb *snail* (*sna*) shadow enhancer ^24^ was placed ∼7.5 kb downstream of the promoter region, which is similar to the enhancer-promoter distance at the endogenous *sna* locus. Transcription of the *MS2-yellow* is dependent on activation signal from linked enhancer since its expression is entirely abolished upon deletion of enhancer sequence from the locus ^25^. For simultaneous visualization of intergenic transcription at the enhancer region, a sequence cassette containing 24x PP7 repeats was fused with the enhancer (Figure 1A; top). CAGE-seq data shows that basal level of non-coding transcription at the *sna* shadow enhancer is below the detection limit (Figure S1A). Consistent with this, enhancer-derived transcripts were not detected at the region where *sna* gene is actively transcribed by fluorescence *in situ* hybridization (FISH) (Figure S1C and D). We then examined if non-coding enhancer transcription can be triggered from this synthetic locus. A short 155-bp DNA fragment containing minimal core promoter motifs were placed adjacent to the enhancer to mimic genomic configuration around the transcription start sites (TSSs) of unidirectionally transcribed enhancers (Figure 1A; middle) ^26, 27^. This engineered synthetic locus was then integrated into the same genomic landing site via phiC31-mediated transgenesis ^28^. Notably, this minimal modification successfully triggered production of non-coding transcripts from the intergenic enhancer region at the *sna* expression domain (Figure S1E and F). It has been previously shown that bidirectional elongation complexes seen at mammalian TSSs are less prevalent in *Drosophila* ^29^. Consistent with this, inversion of engineered TSS eliminated transcription from the enhancer region (Figure S1G and H), demonstrating that non-coding intergenic transcription seen in this study is actually unidirectional. Thus, the use of this system enables us to precisely modulate the mode of intergenic transcription, and quantitatively visualize its functional impact on gene activity in developing embryos.

**Figure 1.**
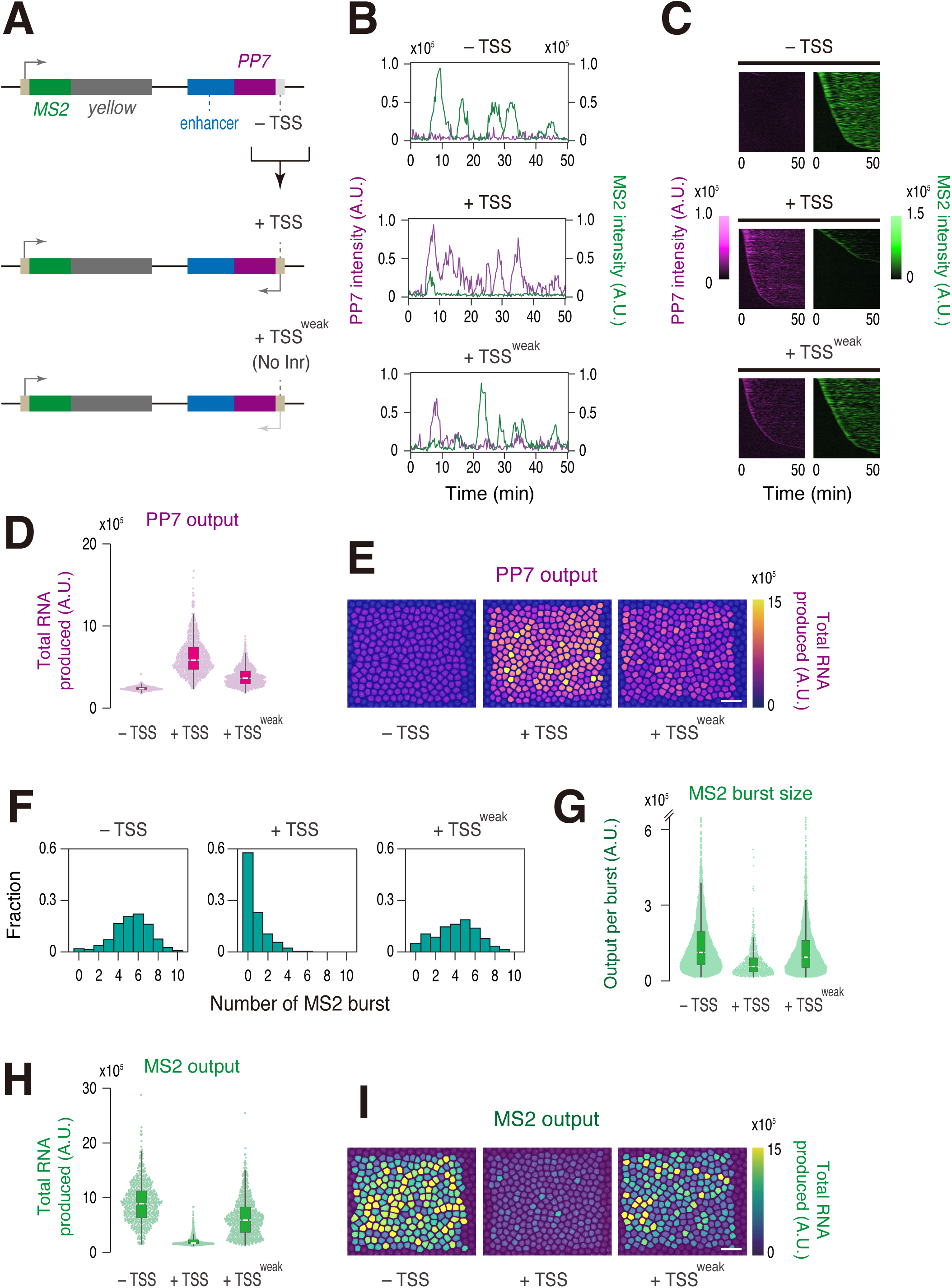
Non-coding enhancer transcription suppresses burst induction. (A) The *yellow* reporter gene containing the *Drosophila* synthetic core promoter (DSCP) and 24x MS2 repeats were placed under the control of the *sna* shadow enhancer fused with 24x PP7 repeats (top). Minimal core promoter motifs were placed adjacent to the enhancer to drive intergenic non-coding transcription (middle). Inr motif at the intergenic TSS was specifically mutated (bottom). (B) Representative trajectories of transcription activities of the reporter locus containing – TSS (top), + TSS (middle), or + TSS^weak^ (Inr mutant; bottom) at the enhancer region. A.U.; arbitrary unit. (C) MS2 and PP7 trajectories for all analyzed nuclei. Each row represents the MS2 or PP7 trajectory for a single nucleus. A total of 676, 719, and 700 ventral-most nuclei, respectively, were analyzed from three independent embryos for the reporter locus containing – TSS (top), + TSS (middle), or + TSS^weak^ (bottom) at the enhancer region. Nuclei were ordered by the onset of MS2 or PP7 transcription in nc14, separately. The same number of nuclei were analyzed hereafter. (D) Boxplot showing the distribution of total output of PP7 transcription. The box indicates the lower (25%) and upper (75%) quantile and the white line indicates the median. Whiskers extend to the most extreme, non-outlier data points. (E) Each nucleus was colored with respect to the total output of PP7 transcription in the representative embryos. The maximum projected image of His2Av-eBFP2 is shown in gray. The image is oriented with anterior to the left and ventral view facing up. Scale bar indicates 20 μm. (F) Histograms showing the distribution of MS2 burst frequency. (G) Boxplot showing the distribution of MS2 burst size. The box indicates the lower (25%) and upper (75%) quantile and the white line indicates the median. Whiskers extend to the most extreme, non-outlier data points. A total of 3607, 536, and 2845 MS2 bursts, respectively, were analyzed for the reporter locus containing – TSS, + TSS, or + TSS^weak^ at the enhancer region. The double hash mark on the y-axis indicates that >99% of the data points are presented. (H) Boxplot showing the distribution of total output of MS2 transcription. The box indicates the lower (25%) and upper (75%) quantile and the white line indicates the median. Whiskers extend to the most extreme, non-outlier data points. (I) Each nucleus was colored with respect to the total output of MS2 transcription in the representative embryos. The maximum projected image of His2Av-eBFP2 is shown in gray. The image is oriented with anterior to the left and ventral view facing up. Scale bar indicates 20 μm.

### Non-coding enhancer transcription attenuates expression of linked gene

We next sought to determine how the emergence of non-coding enhancer transcription impacts expression profiles of linked MS2 reporter gene in living embryos. Nascent RNA production from the MS2 reporter gene and PP7-tagged enhancer was simultaneously visualized with maternally provided MCP-GFP and mCherry-PCP fusion proteins from the entry into nuclear cycle 14 (nc14) (Movie S1). Consistent with FISH analysis (Figure S1C-F), entire population of the ventral-most nuclei produced clear PP7 signal when TSS was engineered at the intergenic region (Figure 1B-E; – TSS vs. + TSS). We then analyzed expression profiles of the MS2 reporter gene in the same embryos. Unexpectedly, we observed a sharp reduction in the level of MS2 activation when transcription was triggered at the enhancer region (Figure 1B and C). Heatmap analysis revealed that there is an overall delay in activating MS2 reporter transcription in the presence of enhancer transcription (Figure 1C). Indeed, ∼60% of ventral-most nuclei never experienced MS2 transcription during the analysis while non-coding enhancer transcription occurred much more ubiquitously and rapidly (Figure S2A; middle). Consistent with this, there was a dramatic decrease in the frequency of MS2 burst (Figure 1F). In addition, the size of individual MS2 burst (i.e., number of nascent transcripts produced per burst) became ∼2-fold smaller upon induction of enhancer transcription (Figure 1G). As a consequence, total output of MS2 transcription was reduced by > 80% (Figure 1H and I). Same results were also seen when the *sna* shadow enhancer was replaced with another well-characterized developmental enhancer, *rhomboid* neuroectoderm element (*rho* NEE) ^30^ (Figure S1B and S3). These results together give rise to the possibility that non-coding enhancer transcription somehow attenuates induction of transcriptional bursting from linked gene in developing embryos.

It has been previously shown that Initiator (Inr) motif is enriched around the TSSs of transcribed enhancers ^e.g., 26, 27, 31, 32^. Intergenic TSS used in this study also contains Inr motif. To examine if an extent of enhancer transcription correlates with its inhibitory function, we next produced a synthetic locus where Inr motif at the intergenic TSS was specifically mutated (Figure 1A; bottom). As expected, the level of enhancer transcription was significantly reduced upon Inr mutation (Figure 1B-E; + TSS^weak^). Conversely, burst profile and transcriptional output of the MS2 reporter gene were largely restored in this genomic configuration (Figure 1F-I). Overall, these results are consistent with the idea that non-coding intergenic transcription can flexibly modulate the functionality of enhancers in an activity-dependent manner.

### Mechanism of transcriptional attenuation by non-coding enhancer transcription

Our data suggest that non-coding intergenic transcription reduces an efficiency of burst induction. To obtain mechanistic insights into this process, we first examined the possibility that non-coding transcripts themselves play a role in attenuating burst induction from the MS2 reporter gene. Indeed, there are several examples where enhancer-derived non-coding transcripts significantly alter gene activity *in trans* ^33, 34^. According to this *trans*-acting model, it is expected that enhancer-derived non-coding transcripts produced from the PP7 allele located on the other homologous chromosome can also impact expression profile of the MS2 allele lacking intergenic TSS *in trans* (Figure 2A). Live-imaging analysis of resulting embryos revealed that the MS2 reporter gene maintains a high level of transcription irrespectively of the induction of non-coding enhancer transcription from the other allele (Figure 2B-D, Movie S2). Consistently, burst profile and total output of the MS2 reporter gene remained to be essentially unchanged (Figure 2E-G). Thus, it is likely that enhancer-derived non-coding transcripts do not possess an ability to modulate gene activity *in trans*.

**Figure 2.**
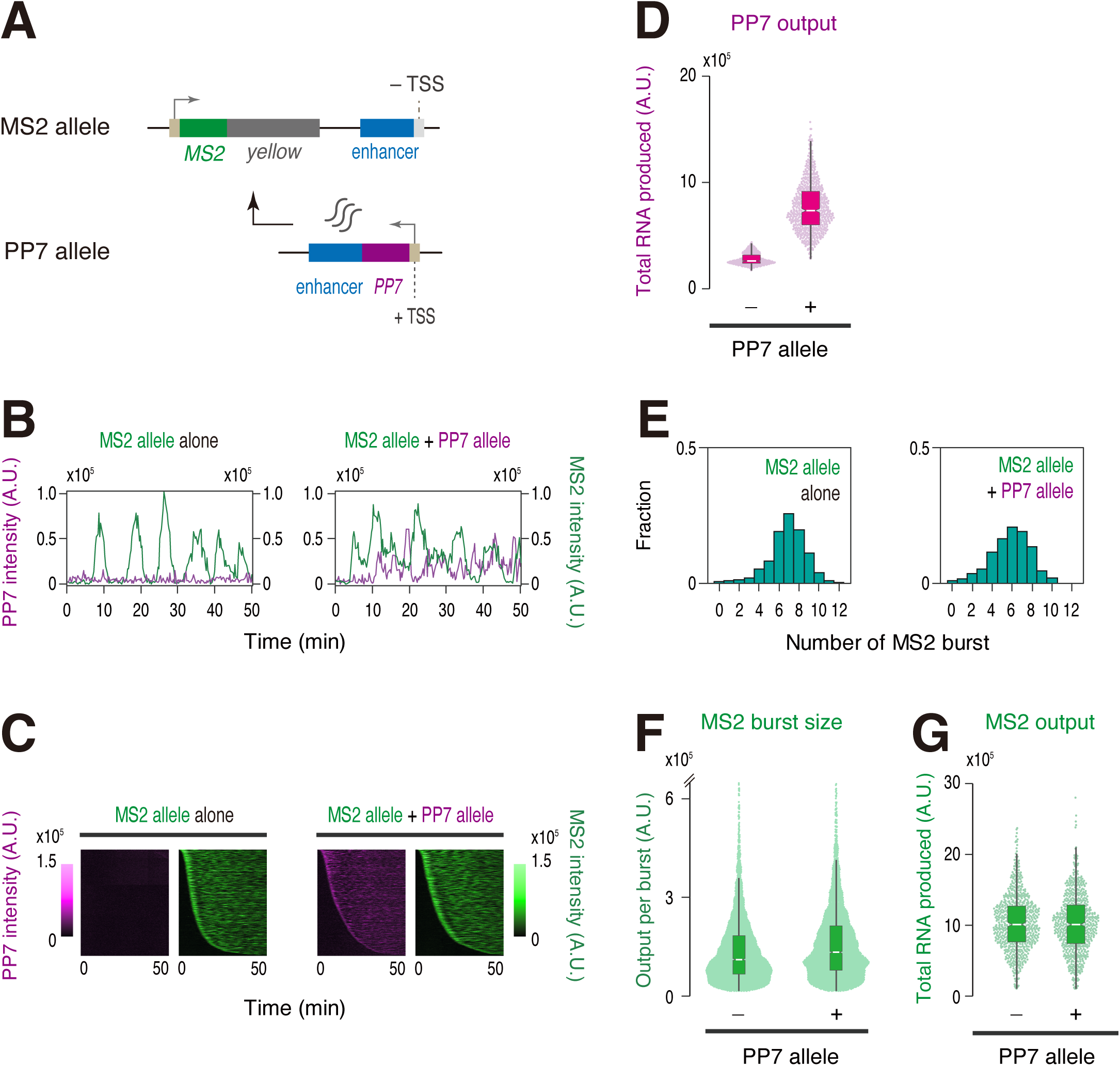
Non-coding transcripts do not affect target gene expression *in trans*. (A) The *MS2-yellow* reporter gene was placed under the control of non-transcribed *sna* shadow enhancer (MS2 allele). Production of enhancer-derived non-coding transcripts was driven from the *sna* shadow enhancer located on the other homologous chromosome (PP7 allele). (B) Representative trajectories of transcription activities of the MS2 allele with (right) or without (left) the PP7 allele. A.U.; arbitrary unit. (C) MS2 and PP7 trajectories for all analyzed nuclei. Each row represents the MS2 or PP7 trajectory for a single nucleus. A total of 719 and 704 ventral-most nuclei, respectively, were analyzed from three independent embryos lacking (left) or containing (right) the PP7 allele. Nuclei were ordered by the onset of MS2 or PP7 transcription in nc14, separately. The same number of nuclei were analyzed hereafter. (D) Boxplot showing the distribution of total output of PP7 transcription. The box indicates the lower (25%) and upper (75%) quantile and the white line indicates the median. Whiskers extend to the most extreme, non-outlier data points. (E) Histograms showing the distribution of MS2 burst frequency. (F) Boxplot showing the distribution of MS2 burst size. The box indicates the lower (25%) and upper (75%) quantile and the white line indicates the median. Whiskers extend to the most extreme, non-outlier data points. A total of 4917 and 4109 MS2 bursts, respectively, were analyzed for embryos lacking or containing the PP7 allele. The double hash mark on the y-axis indicates that >99% of the data points are presented. (G) Boxplot showing the distribution of total output of MS2 transcription. The box indicates the lower (25%) and upper (75%) quantile and the white line indicates the median. Whiskers extend to the most extreme, non-outlier data points.

We then explored the possibility that convergent transcription from the enhancer region counteracts target gene activation by allowing enhancer-transcribing Pol II to penetrate into the neighboring MS2 gene body. In our system, both the MS2 and PP7 transcription units were designed to contain termination site at the 3’ end. In yeast, it has been reported that head-to-head collision of Pol II interrupts expression of convergently transcribed genes ^35^. To test if a similar mechanism also takes place in *Drosophila*, we produced a reporter strain that contains transcribing enhancer in a tandem orientation (Figure 3A, Movie S3). Equivalently strong enhancer transcription was seen in either genomic configuration (Figure 3B-D). Importantly, we found that attenuation of MS2 reporter transcription is only partially relieved in a tandem orientation (Figure 3E-G). Indeed, total output of MS2 transcription remained to be ∼70% lower comparing to the reporter locus lacking intergenic TSS (Figure S2B). These results suggest that transcriptional readthrough or Pol II collision is not a major source of transcriptional attenuation seen in this study. In this regard, our mechanism is clearly different from transcriptional attenuation of host gene expression by intragenic enhancers in mammalian system where Pol II collision is suggested to play a central role ^36^.

**Figure 3.**
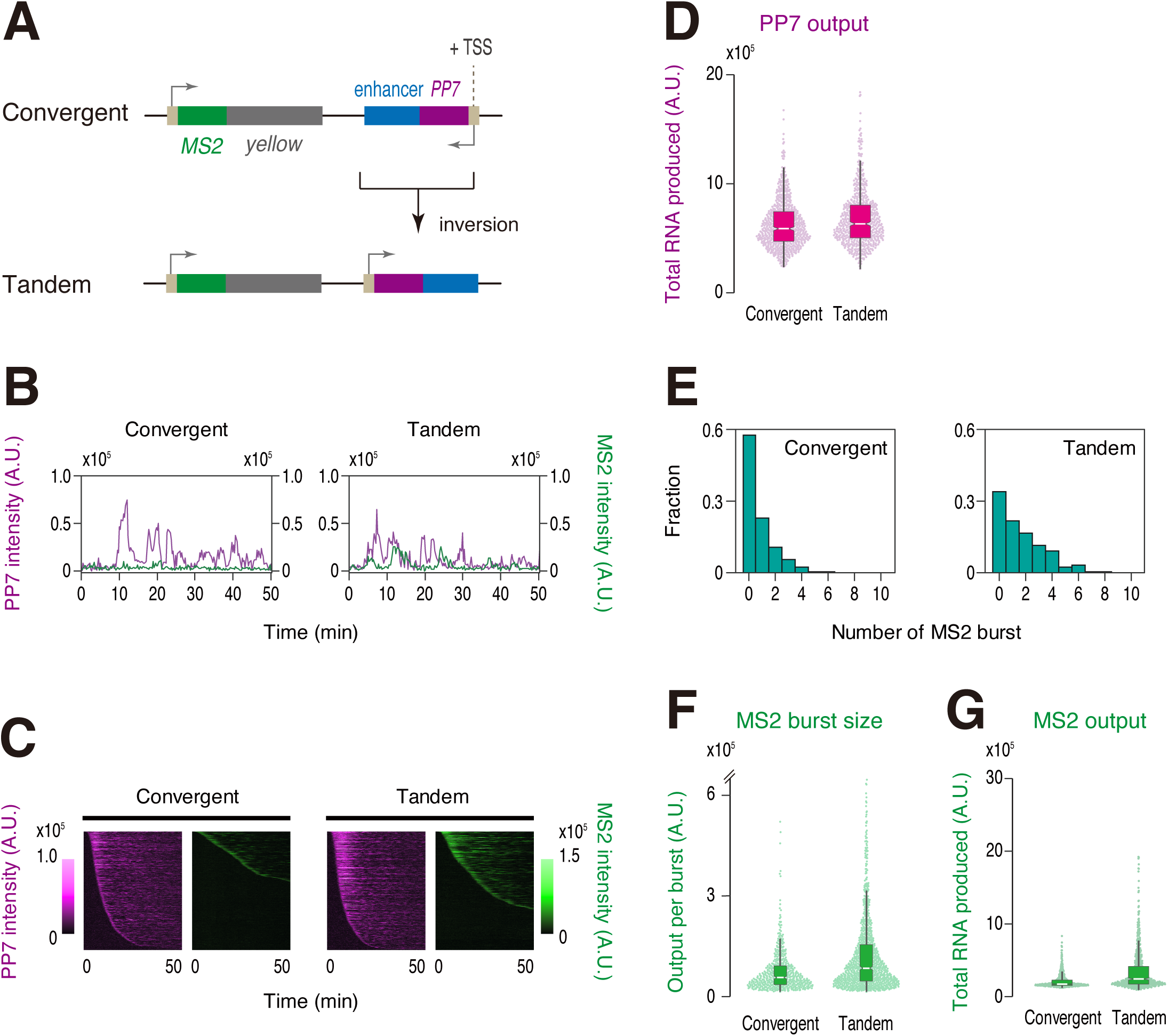
Non-coding enhancer transcription in a tandem orientation also suppresses burst induction. (A) Transcribing *sna* shadow enhancer was placed in a tandem orientation relative to the *MS2-yellow* reporter gene. (B) Representative trajectories of transcription activities of the reporter locus driving non-coding enhancer transcription in a convergent (left) or a tandem orientation (right). A.U.; arbitrary unit. (C) MS2 and PP7 trajectories for all analyzed nuclei. Each row represents the MS2 or PP7 trajectory for a single nucleus. A total of 719 and 638 ventral-most nuclei, respectively, were analyzed from three independent embryos for the reporter locus driving non-coding enhancer transcription in a convergent (left) or a tandem orientation (right). Nuclei were ordered by the onset of MS2 or PP7 transcription in nc14, separately. The same number of nuclei were analyzed hereafter. Panel of Convergent is the same as the panel of + TSS shown in Figure 1C. (D) Boxplot showing the distribution of total output of PP7 transcription. The box indicates the lower (25%) and upper (75%) quantile and the white line indicates the median. Whiskers extend to the most extreme, non-outlier data points. Plot of Convergent is the same as the plot of + TSS shown in Figure 1D. (E) Histograms showing the distribution of MS2 burst frequency. Plot of Convergent is the same as the plot of + TSS shown in Figure 1F. (F) Boxplot showing the distribution of MS2 burst size. The box indicates the lower (25%) and upper (75%) quantile and the white line indicates the median. Whiskers extend to the most extreme, non-outlier data points. A total of 536 and 1055 MS2 bursts, respectively, were analyzed for the reporter locus driving non-coding enhancer transcription in a convergent or tandem orientation. The double hash mark on the y-axis indicates that >99% of the data points are presented. Plot of Convergent is the same as the plot of + TSS shown in Figure 1G. (G) Boxplot showing the distribution of total output of MS2 transcription. The box indicates the lower (25%) and upper (75%) quantile and the white line indicates the median. Whiskers extend to the most extreme, non-outlier data points. Plot of Convergent is the same as the plot of + TSS shown in Figure 1H.

Previous studies suggested that two linked transcription units can compete with each other for an interaction with a shared enhancer when they are placed under the control of a single enhancer ^e.g., 37,38,39^. More recently, it has also been suggested that high levels of nascent RNA act as a negative feedback mechanism to downregulate gene transcription^20^. To test if transcriptional attenuation is caused by promoter competition or RNA-mediated feedback mechanism, we produced a synthetic locus where PP7 transcription is driven in an outward orientation relative to the enhancer (Figure 4A, Movie S4). In this reporter locus, we observed even stronger PP7 transcription comparing to the original reporter configuration (Figure 4B-D). According to promoter competition or RNA-mediated feedback hypothesis, it is expected that increased PP7 activity will lead to further diminishment of target gene activity. However, contrary to this expectation, burst induction from the MS2 reporter gene was largely restored upon inversion of PP7 transcription (Figure 4E-G, Figure S3), suggesting that neither promoter competition nor RNA-mediated feedback mechanism can account for the attenuation of target gene expression seen in this study. Instead, our results strongly indicate that Pol II elongation across the internal core region of enhancer is responsible for limiting their capability of inducing transcriptional bursting from linked gene.

**Figure 4.**
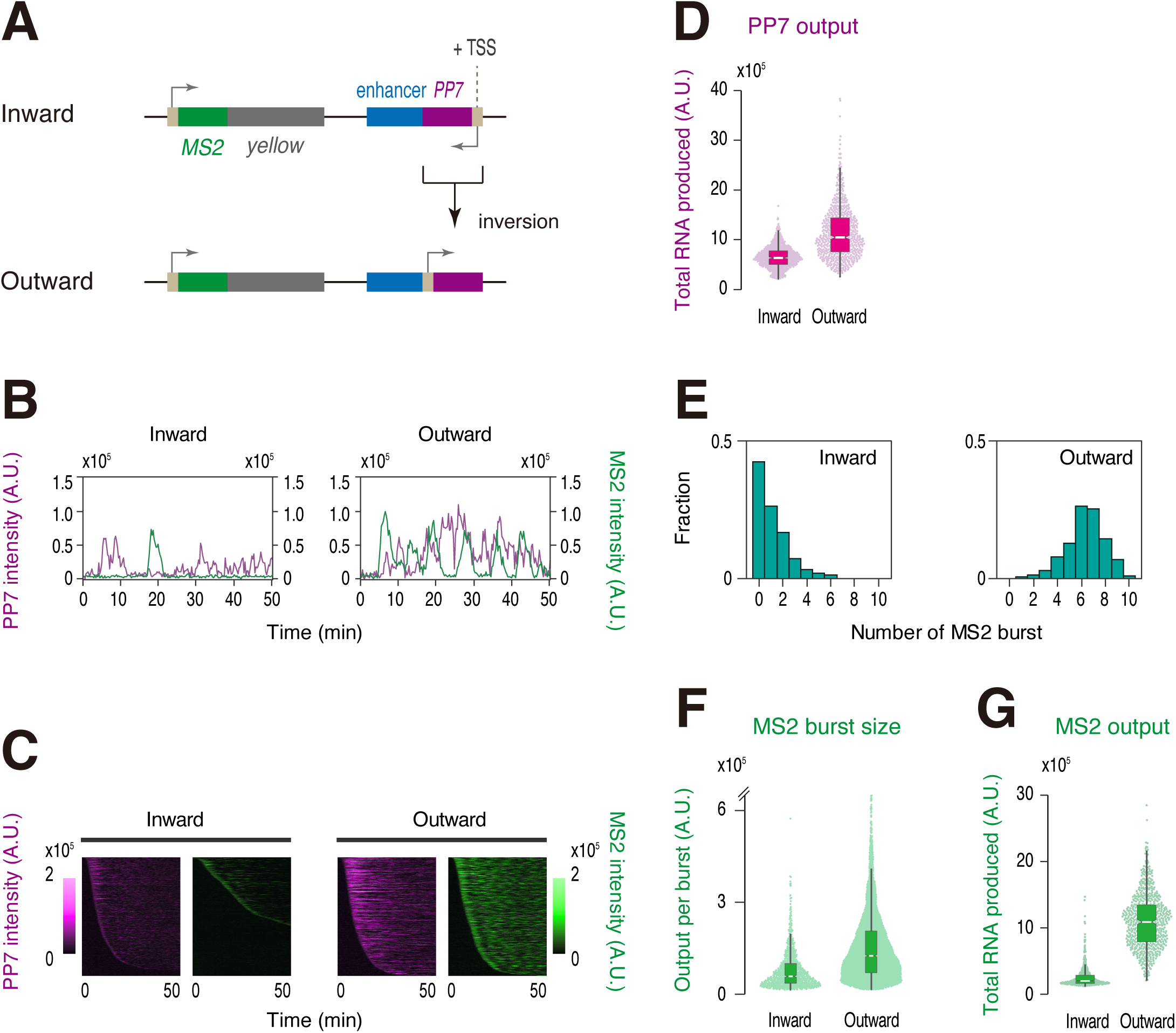
Enhancer self-transcription is required for target gene attenuation. (A) PP7 transcription unit at the enhancer region was inverted to drive non-coding transcription in an outward orientation relative to the *sna* shadow enhancer. (B) Representative trajectories of transcription activities of the reporter locus driving PP7 transcription in an inward (left) or an outward orientation (right). A.U.; arbitrary unit. (C) MS2 and PP7 trajectories for all analyzed nuclei. Each row represents the MS2 or PP7 trajectory for a single nucleus. A total of 739 and 666 ventral-most nuclei, respectively, were analyzed from three independent embryos for the reporter locus driving PP7 transcription in an inward (left) or an outward orientation (right). Nuclei were ordered by the onset of MS2 or PP7 transcription in nc14, separately. The same number of nuclei were analyzed hereafter. (D) Boxplot showing the distribution of total output of PP7 transcription. The box indicates the lower (25%) and upper (75%) quantile and the white line indicates the median. Whiskers extend to the most extreme, non-outlier data points. (E) Histograms showing the distribution of MS2 burst frequency. (F) Boxplot showing the distribution of MS2 burst size. The box indicates the lower (25%) and upper (75%) quantile and the white line indicates the median. Whiskers extend to the most extreme, non-outlier data points. A total of 845 and 4220 MS2 bursts, respectively, were analyzed for the reporter locus driving PP7 transcription in an inward or an outward orientation. The double hash mark on the y-axis indicates that >99% of the data points are presented. (G) Boxplot showing the distribution of total output of MS2 transcription. The box indicates the lower (25%) and upper (75%) quantile and the white line indicates the median. Whiskers extend to the most extreme, non-outlier data points.

### Modulation of hub formation via non-coding enhancer transcription

Then, how does Pol II elongation across the internal core enhancer region lead to attenuation of transcriptional bursting at the molecular level? Notably, recent cryo-EM studies suggested that elongating Pol II can progressively peel off template DNA from histone surfaces ^40–42^. In analogy to this, we hypothesized that elongating Pol II facilitates eviction of transcription factors from the enhancer region (Figure 5A). As seen in ChIP-seq profile (Figure S1A), the *sna* shadow enhancer used in this study is enriched with a sequence-specific transcription factor Dorsal (Dl), a homolog of mammalian NF-κB. To directly visualize how enhancer self-transcription impacts nuclear localization of Dl, we employed CRISPR/Cas9 genome-editing to fuse GFP to the C-terminus of the protein. It was confirmed that resulting Dl-GFP strain is homozygous viable and fertile. We then monitored endogenous Dl together with PP7 signal using Zeiss Airyscan2 super-resolution imaging system (Figure 5B). We first analyzed a synthetic locus where non-inhibitory PP7 transcription occurs in an outward orientation relative to the enhancer (Figure 4A; bottom). Following previous imaging studies ^43, 44^, we obtained averaged profile of Dl signal surrounding active PP7 signal over 800 nuclei from 50 independent embryos at nc14 (Movie S5 and S6). We observed a sharp enrichment of Dl signal at the site of transcription comparing to randomly selected locations within nuclei (Figure 5C and D; left). This is consistent with a recent imaging study showing that Dl forms a cluster or “hub” at the site of transcription in early embryos ^45^. We then analyzed the formation of Dl hub at the reporter locus where inhibitory PP7 transcription is driven at the enhancer region (Figure 4A; top). There was a clear reduction in the local concentration of Dl in this experimental setup (Figure 5C and D; right), suggesting that enhancer self-transcription acts to interrupt stable association of transcription factors at the regulatory region. To further test this idea, we next examined nuclear distribution of a pioneering transcription factor Zelda (Zld) since it was recently reported that Dl cooperates with Zld to form a hub ^45^. Visualization of endogenous Zld-GFP revealed that it also forms a hub at the enhancer region (Figure S4A and B; left). As seen for Dl, formation of Zld hub became much less clear in the presence of inhibitory PP7 transcription (Figure S4A and B; right), supporting the idea that enhancer transcription exerts its negative regulatory function through the modulation of hub formation. Importantly, we noticed that decrease in the local concentration of Dl and Zld was not as drastic as reduction in gene activity we observed (Figure 4). Taking into account recent studies showing that subtle changes in enhancer-promoter interactions can create large changes in transcriptional output ^46, 47^, changes in the size of transcription hub may also have non-linear impact on gene activity. It is also conceivable that Pol II elongation further impairs functional integrity of activator hub formed at the enhancer region by interrupting subsequent recruitment of other key co-factors (see Discussion). Overall, our results are consistent with the idea that intergenic non-coding transcription modulates enhancer activity by impacting the formation and the function of transcription hub in developing embryos.

**Figure 5.**
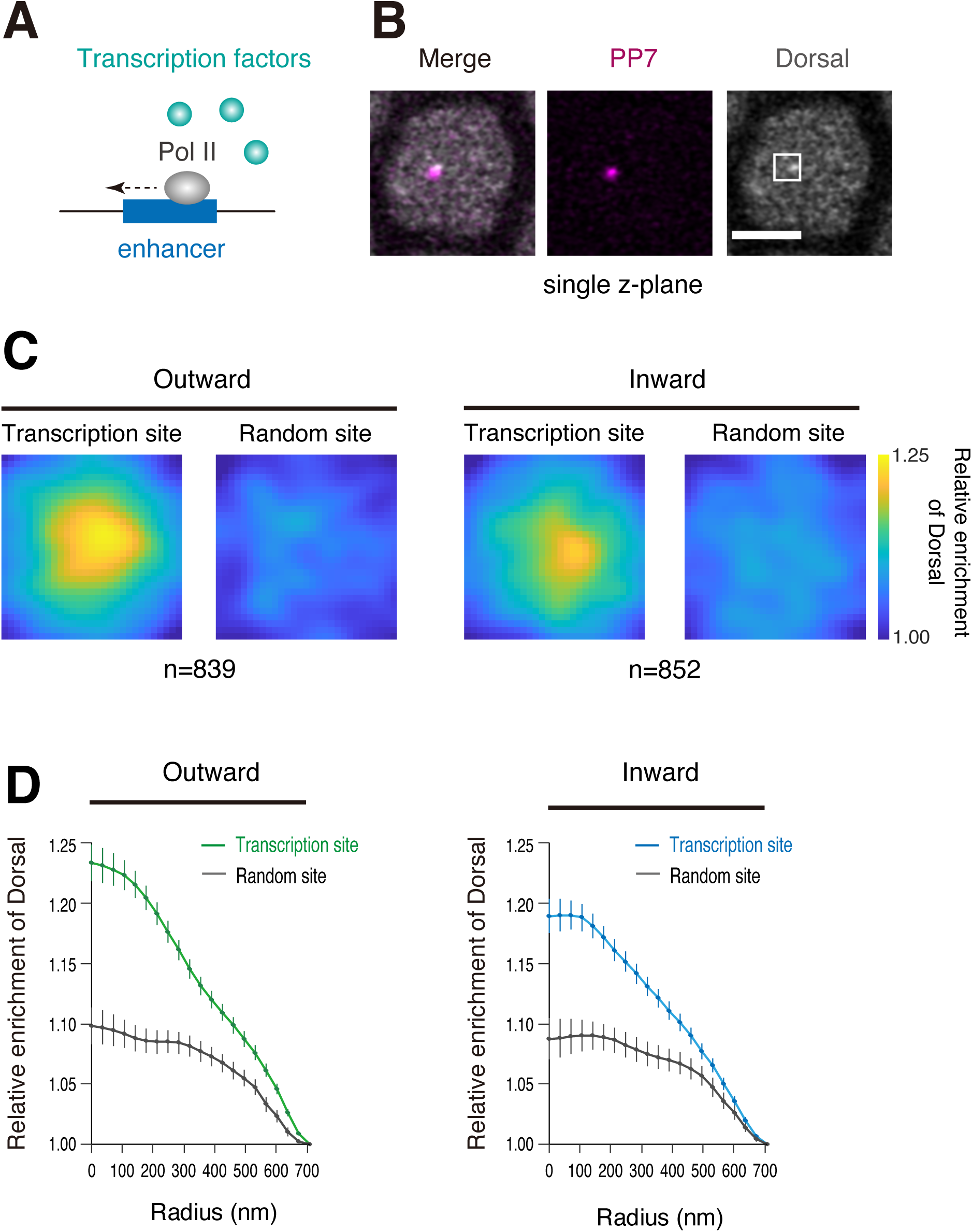
Enhancer self-transcription decreases local concentration of Dl activator. (A) Elongating Pol II mediates eviction of transcription factors from the enhancer region. (B) Representative Airyscan images of Dorsal-GFP and PP7 transcription dot in the single z-plane containing the brightest PP7 signal. White square indicates analyzed region (29 x 29 pixels) centering the brightest PP7 signal. Scale bar indicates 3 μm. (C) Heatmaps showing the averaged distribution of Dorsal-GFP centering the PP7 transcription site or random site. A total of 839 and 852 PP7-transcribing nuclei, respectively, were obtained from 50 independent embryos for the reporter locus driving PP7 transcription in an outward (left) or an inward orientation (right). (D) Radial profiles of the averaged Dorsal-GFP distribution shown in (C). Error bar represents standard error of the mean.

### Naturally transcribing *Ubx* enhancer can effectively activate gene transcription

Our experiments so far focused on the analysis of synthetically engineered enhancers. We next sought to visualize activities of naturally transcribing enhancers during early embryogenesis. For this purpose, we first analyzed publicly available 2-4h CAGE-seq dataset ^48^ to quantify read counts for >650 functionally verified enhancers in fly embryos ^49^, resulting in the identification of highly transcribing enhancers at this developmental stage (Figure 6A). We found that enhancers regulating the expression of key developmental genes such as Hox genes *Ultrabithorax* (*Ubx*) and *Abdominal-B* (*Abd-B*) represent one of the most actively transcribing enhancers (Figure 6A, Table 1). Among these, we were particularly interested in the *Ubx* enhancer since this region corresponds to the classical *bx* region enhancer (BRE) that is known to be required for *Ubx* expression at the middle part of embryos (Figure 6B; bottom) ^50^ and for the development of metathoracic segment in adult flies ^51^. A sharp and selective CAGE-seq peak was seen for one specific DNA strand at this region (Figure 6B; top), indicating that BRE undergoes unidirectional non-coding transcription in early embryos. Intriguingly, we noticed that unidirectional transcription initiated from BRE is facing toward the opposite orientation relative to the transcription factor binding sites within the enhancer (Figure 6B), giving rise to a possibility that BRE is naturally optimized to drive non-coding transcription in a manner that does not impede functions of bound transcription factors. To experimentally test this idea, PP7-tagged BRE was linked to the MS2 reporter and its activity was visualized in living embryos (Figure 6C; top). Consistent with CAGE-seq profile (Figure 6B), we reproducibly observed a clear PP7 signal originating from BRE (Figure 6D and E, Movie S7). Importantly, BRE was found to drive a high level of transcription from the MS2 reporter gene (Figure 6D and E), supporting the idea that BRE accommodates non-coding enhancer transcription in a manner that does not impede activation of linked gene. We next divided nuclei into two groups according to the presence or absence of PP7 transcription, and compared MS2 profiles between them. Intriguingly, BRE was found to more effectively activate linked MS2 reporter gene in the presence of PP7 transcription (Figure 6F, Figure S5), implicating that natural configuration of BRE permits non-coding enhancer transcription to exert positive regulatory function. We next engineered this synthetic locus to invert the orientation of BRE-PP7 cassette (Figure 6C; middle). As expected, PP7 signal was lost in this setup (Figure 6D and E; inverted BRE). In addition, we observed a partial reduction in the level of MS2 activity (Figure 6D and E; inverted BRE), which is consistent with our preceding result showing that non-coding transcription in a convergent orientation leads to minor suppression of gene activity (Figure 3). We then engineered BRE to contain additional intergenic TSS to foster Pol II elongation across the transcription factor binding sites within the enhancer (Figure 6C; bottom). Intriguingly, this led to dramatic reduction in the efficiency of burst induction (Figure 6D, E, G-I; additional TSS) and the level of total RNA production from the MS2 reporter gene (Figure 6J). Overall, these results are consistent with the idea that the natural configuration of BRE enables cells to effectively activate gene transcription together with non-coding transcription at the same time. Intriguingly, similar configuration was seen for other transcribing enhancers identified in this study (Figure S6). Thus, it is tempting to speculate that configurations of non-coding TSSs have been evolutionarily defined, in part, by their orientation relative to the nearby transcription factor binding sites and accompanying regulatory effects at each genomic locus (see Discussion).

**Figure 6.**
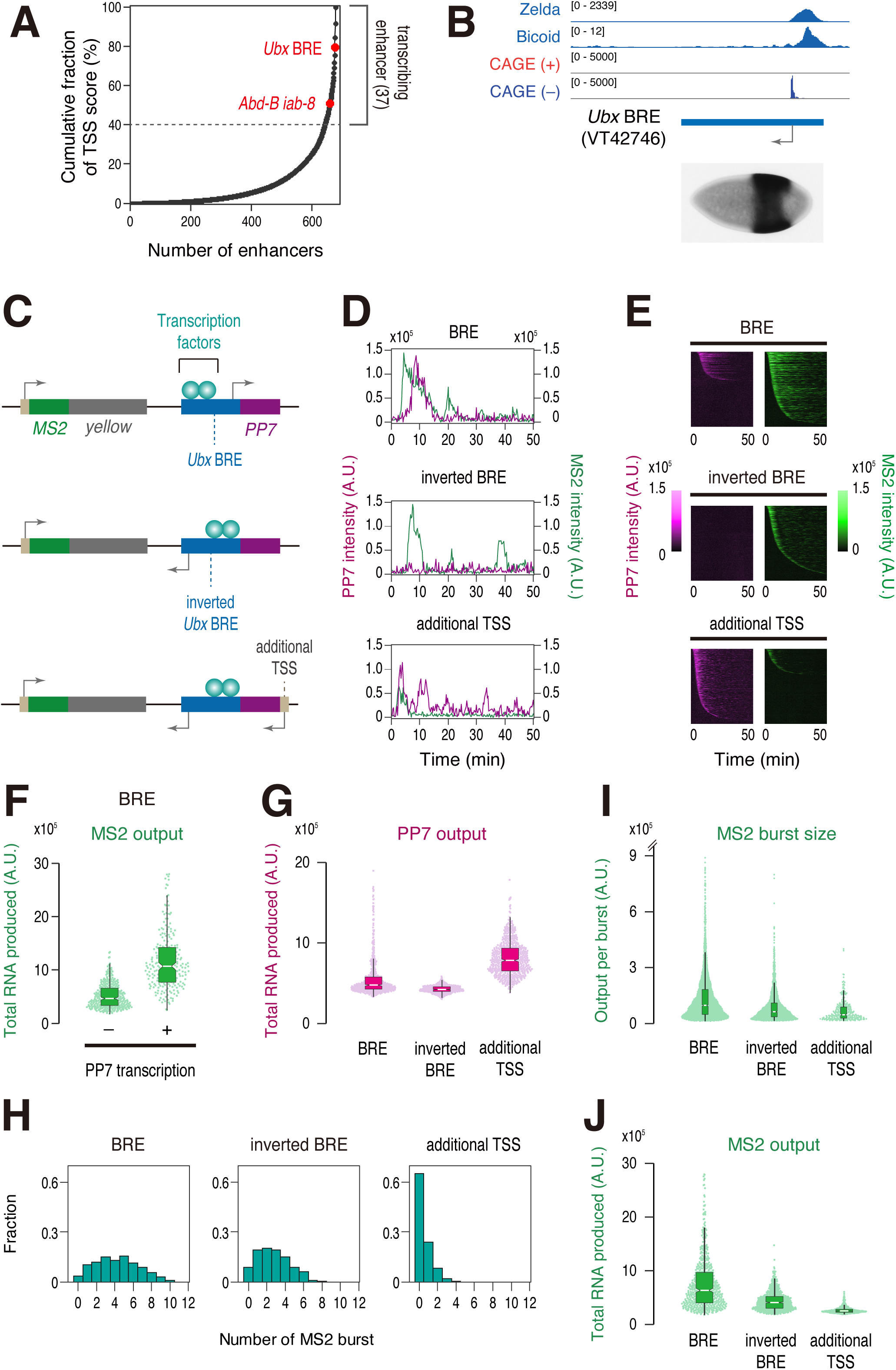
Natural *Ubx* BRE co-activates non-coding transcription and gene transcription effectively. (A) Cumulative fraction of TSS score for all the analyzed developmental enhancers. A total of 678 functionally validated enhancers was analyzed ^60^. The dashed line represents the threshold for determining transcribed enhancers at early developmental stage. (B) Organization of the endogenous *Ubx* BRE. Zelda ChIP-seq data from nc13 WT embryos (GSM763061) ^61^, Bicoid-GFP ChIP-seq from nc14 WT embryos (GSE86966) ^62^, integrated CAGE-seq data on positive and negative strand from 2-to 4-h WT embryos (ERR1425056) ^58^ were visualized with Integrative Genomics Viewer. A picture of an early embryo showing the activity of *Ubx* BRE (VT42746; ∼2.2 kb in length) was taken from Fly Enhancers ^60^. (C) A ∼1.2-kb DNA fragment containing *Ubx* BRE was fused with 24x PP7 repeats, and placed downstream of the MS2 reporter gene (top). The orientation of BRE-PP7 cassette was inverted (middle). Minimal core promoter motifs were placed adjacent to the enhancer to drive additional non-coding enhancer transcription (bottom). (D) Representative trajectories of transcription activities of the reporter locus containing BRE (top), inverted BRE (middle), or inverted BRE with additional TSS (bottom). A.U.; arbitrary unit. (E) MS2 and PP7 trajectories for all analyzed nuclei. Each row represents the MS2 or PP7 trajectory for a single nucleus. A total of 582, 654, and 636 nuclei, respectively, were analyzed from three independent embryos for the reporter locus containing BRE (top), inverted BRE (middle), or inverted BRE with additional TSS (bottom). Nuclei were ordered by the onset of MS2 or PP7 transcription in nc14, separately. The same number of nuclei were analyzed hereafter. (F) Boxplot showing the distribution of total output of MS2 transcription. A total of 359 and 223 nuclei, respectively, were analyzed for those did not or did undergo PP7 transcription from BRE. The box indicates the lower (25%) and upper (75%) quantile and the white line indicates the median. Whiskers extend to the most extreme, non-outlier data points. (G) Boxplot showing the distribution of total output of PP7 transcription. The box indicates the lower (25%) and upper (75%) quantile and the white line indicates the median. Whiskers extend to the most extreme, non-outlier data points. (H) Histograms showing the distribution of MS2 burst frequency. (I) Boxplot showing the distribution of MS2 burst size. The box indicates the lower (25%) and upper (75%) quantile and the white line indicates the median. Whiskers extend to the most extreme, non-outlier data points. A total of 2473, 1792, and 310 MS2 bursts, respectively, were analyzed for the reporter locus containing BRE, inverted BRE, or inverted BRE with additional TSS. The double hash mark on the y-axis indicates that >99% of the data points are presented. (J) Boxplot showing the distribution of total output of MS2 transcription. The box indicates the lower (25%) and upper (75%) quantile and the white line indicates the median. Whiskers extend to the most extreme, non-outlier data points.

## Discussion

In this study, we have successfully established a live-imaging system that permits simultaneous visualization of non-coding enhancer transcription together with gene transcription at the single-cell resolution in *Drosophila*. By combining a series of genome-engineering and genetic approaches, we have provided several lines of evidence that intergenic non-coding transcription can flexibly modulate enhancer function in an orientation- and activity-dependent manner (Figure 1-4). Super-resolution imaging and genome-editing analysis further demonstrated that enhancer self-transcription impacts molecular crowding of transcription factors and hub formation at the gene locus (Figure 5, Figure S4). We speculate that elongating Pol II acts as a steric hindrance for stable association of transcription factors with their cognate enhancers, which may further interrupt subsequent recruitment of co-activators and other key transcriptional apparatus to the gene locus. Consistent with our results, it has been previously reported that non-coding transcription at the intergenic regulatory region suppresses expression of *SER3* gene by inducing dissociation of transcription factors from their binding sites in yeast ^18, 19^. Transcription of yeast *ADH1* gene is also shown to be suppressed through a similar mechanism during zinc starvation ^15^. Our data together with these preceding studies implicate that attenuation of gene activity by non-coding enhancer transcription is an ancient mechanism conserved from fungi to arthropods. It might be possible that cells utilize this system to dynamically modulate the level of gene activity by changing the directionality and/or strength of intergenic non-coding transcription in response to intrinsic and extrinsic signals such as developmental timing and nutrient condition under natural environment. Acquisition of novel intergenic TSSs may also have contributed to diversification of enhancer function during the process of animal evolution as we were able to produce a broad spectrum of regulatory activities from the same enhancer just by engineering the mode of intergenic non-coding transcription. As exemplified in the analysis of *Ubx* BRE (Figure 6), some enhancers appear to preferentially contain non-inhibitory TSSs facing toward the opposite orientation relative to the transcription factor binding sites. Under its natural enhancer configuration, non-coding transcription from BRE was found to positively correlate with bursting activities of linked gene (Figure 6F, Figure S5), giving rise to the possibility that there are a class of enhancers that utilize non-coding transcription to facilitate gene expression. Our super-resolution imaging analysis indicates that enhancer transcription exerts its negative regulatory function by interrupting the formation and the function of transcription hub at the gene locus (Figure 5, Figure S4). Taking this into consideration, it is also conceivable that enhancer transcription exerts its positive regulatory function by facilitating release of enhancer-bound repressors and/or inhibitory nucleosomes in certain circumstances. Importantly, our data showed that positive effects of BRE non-coding transcription can be easily lost by inverting its orientation or placing additional intergenic TSS (Figure 6), suggesting that regulatory outcome of enhancer transcription is highly dependent on its surrounding genomic context. Thus, we speculate that mode of enhancer transcription has been evolutionally defined, in part, by the balance between its negative and positive regulatory effects at each genomic locus. This “mixed effect model” may also explain the reason why there is only a weak correlation (*ρ* = 0.24) between the level of non-coding transcription and the enhancer strength in *Drosophila* ^32^. Clearly, future studies are needed to fully elucidate biological roles of enhancer transcription by visualizing its regulatory function at the context of endogenous genome. We believe that our study will serve as a critical starting point toward full understanding of multifaced functions of non-coding enhancer transcription in the control of gene expression during animal development.

## Acknowledgement

We thank Koji Kawasaki for helping analysis of Airyscan imaging data, Hitomi Takishita and Misako Sato for fly husbandry, and the Bloomington *Drosophila* Stock Center for fly strains. We are also grateful to members of the Fukaya laboratory for critical comments on the manuscript. This work was supported by JST FOREST program (JPMJFR214W), the Grant-in-Aid for Scientific Research (B) (22H02544), the Grant-in-Aid for Scientific Research on Innovative Areas (Research in a Proposed Research Area) (22H04665), the Grant-in-Aid for Transformative Research Areas (A) (Research in a Proposed Research Area) (21H05742) from the Japan Society for the Promotion of Science, research grants from the Takeda Science Foundation, the Sumitomo Foundation, the Senri Life Science Foundation and the Mitsubishi Foundation.

## Author contributions

K.H performed the experiments and analyzed the data. T.F wrote the initial draft. K.H and T.F edited the manuscript. All the authors discussed the results and approved the manuscript.

## Materials and Methods

### Experimental model

In all experiments, we studied Drosophila melanogaster embryos at nuclear cycle 14. The following fly lines were used in this study: nanos>NLS-mCherry-PCP-NES, His2Av-eBFP2/CyO; nanos>MCP-GFP ^52^, nanos>NLS-mCherry-PCP-NES, His2Av-eBFP2/CyO ^52^, DSCP-MS2-yellow-sna shadow enhancer-PP7-No TSS (this study), DSCP-MS2-yellow-sna shadow enhancer-PP7-TSS (this study), DSCP-MS2-yellow-sna shadow enhancer-PP7-Inverted TSS (this study), DSCP-MS2-yellow-sna shadow enhancer-PP7-TSS^weak^ (this study), DSCP-MS2-yellow-sna shadow enhancer ^53^, TSS-PP7-sna shadow enhancer (this study), DSCP-MS2-yellow-TSS-PP7-sna shadow enhancer (this study), DSCP-MS2-yellow-sna shadow enhancer-TSS-PP7 (this study), DSCP-MS2-yellow-Ubx BRE-PP7 (this study), DSCP-MS2-yellow-inverted Ubx BRE-PP7 (this study), DSCP-MS2-yellow-inverted Ubx BRE-PP7-TSS (this study), DSCP-MS2-yellow-rho NEE-PP7-No TSS (this study), DSCP-MS2-yellow-rho NEE-PP7-TSS (this study), DSCP-MS2-yellow-rho NEE-TSS-PP7 (this study), dorsal-GFP (this study), zelda-GFP (this study).

### Site-specific transgenesis by phiC31 system

All reporter plasmids were integrated into a unique landing site on the third chromosome using VK00033 strain ^54^. phiC31 was maternally provided using *vas-phiC31* strain ^55^. Microinjection was performed as previously described ^56^. In brief, 0-1 h embryos were collected and dechorionated with bleach. Aligned embryos were dried with silica gel for ∼8 min and covered with FL-100-1000CS silicone oil (Shin-Etsu Silicone). Subsequently, microinjection was performed using FemtoJet (Eppendorf) and DM IL LED inverted microscope (Leica) equipped with M-152 Micromanipulator (Narishige). Injection mixture typically contains ∼900 ng/μl plasmid DNA, 5 mM KCl, 0.1 mM phosphate buffer, pH 6.8. mini-white marker was used for screening.

### CRISPR/Cas9-mediated genome editing

pCFD3 gRNA expression plasmid and pBS-GFP-3xFLAG-3xP3-dsRed donor plasmid were co-injected using *nanos-Cas9/CyO* strain ^57^. Injection mixture contains ∼500 ng/μl pCFD3 gRNA expression plasmid and ∼800 ng/μl pBS-GFP-3xFLAG-3xP3-dsRed donor plasmid. Microinjection was performed as described above. 3xP3-dsRed was used for screening. Resulting *dorsal-GFP* flies were crossed with *y[1] w[67c23] P{y[+mDint2]=Crey}1b; sna[Sco]/CyO* (BDSC# 766) to remove 3xP3-dsRed marker from the locus.

### Preparation of probes for *in situ* hybridization

Antisense RNA probes labeled with digoxigenin (DIG RNA Labeling Mix 10 × conc, Roche) or biotin (Biotin RNA Labeling Mix 10 × conc, Roche) were transcribed using *in vitro* Transcription T7 Kit (Takara). Template DNA for *sna* probe was PCR amplified from genomic DNA using primers (5 ’-CGT AAT ACG ACT CAC TAT AGG GCA GTT GGC TTA ACA GTA CTG-3 ’) and (5 ’-ACC TGT CAC AGC CAC CTC AGC-3 ’). Template DNA for *sna* shadow enhancer probe was PCR amplified from pbphi-TSS-PP7-*sna* shadow enhancer plasmid using primers (5 ’-GCA TTG AGG TGT TTT GTT GGT CAA C-3 ’) and (5 ’-CGT AAT ACG ACT CAC TAT AGG GTA AAT TCC GAT TTT TCT TGT-3 ’).

### Fluorescence *in situ* hybridization

Embryos were dechorionated and fixed in fixation buffer (1 ml of 5x PBS, 4 ml of 37% formaldehyde and 5 ml of Heptane) for ∼25 min at room temperature. Antisense RNA probes labeled with digoxigenin (for *sna* shadow enhancer) and biotin (for *sna* gene) were used. Hybridization was performed at 55°C overnight in hybridization buffer (50% formamide, 5x SSC, 50 μg/ml Heparin, 100 μg/ml salmon sperm DNA, 0.1% Tween-20). Subsequently, embryos were washed with hybridization buffer at 55°C and incubated with Western Blocking Reagent (Roche) at room temperature for ∼2 hours. Then, embryos were incubated with sheep anti-digoxigenin (Roche) and mouse anti-biotin (Invitrogen) primary antibody at 4°C for overnight, followed by incubation with Alexa Fluor 555 donkey anti-sheep (Invitrogen) and Alexa Fluor 488 donkey anti-mouse (Invitrogen) fluorescent secondary antibody at room temperature for ∼2 hours. Embryos were mounted in ProLong Gold Antifade Mountant (Thermo Fisher Scientific). Imaging was performed on a Zeiss LSM 900 confocal microscope. Plan-Apochromat 20x / 0.8 N.A. objective was used. Images were acquired with following settings: 1024 x 1024 pixels, 16-bit depth, 15 z-slices separated by 0.5 μm. Maximum projection was obtained for all z-sections, and resulting image was rotated with bilinear interpolation method. Brightness of images was linearly adjusted using Fiji.

### Preparation of embryos for live-imaging

For MS2/PP7 two-color live-imaging, virgin females of *nanos>NLS-mCherry-PCP-NES, His2Av-eBFP2/CyO*; *nanos>MCP-GFP* were mated with homozygous males carrying the MS2/PP7 reporter allele. In Figure 2, virgin females of *nanos>NLS-mCherry-PCP-NES, His2Av-eBFP2/CyO*; *nanos>MCP-GFP* were first crossed with homozygous males carrying the PP7 allele. Virgin females of resulting trans-heterozygote were then mated with homozygous males carrying the MS2 allele. For Airyscan imaging, virgin females of *nanos>NLS-mCherry-PCP-NES, His2Av-eBFP2/CyO* were first crossed with homozygous males of *dorsal-GFP* or *zelda-GFP*. Virgin females of resulting trans-heterozygote were then mated with homozygous males carrying the MS2/PP7 reporter allele. The resulting embryos were dechorionated and mounted between a polyethylene membrane (Ube Film) and a coverslip (18 mm x 18 mm), and embedded in FL-100-450CS (Shin-Etsu Silicone).

### MS2/PP7 two-color live-imaging

Embryos were imaged using a Zeiss LSM 900 confocal microscope. Plan-Apochromat 40x / 1.4 N.A. oil immersion objective was used. Images were acquired with following settings: 512 x 512 pixels, 16-bit depth, 18 z-slices separated by 0.6 μm, ∼16.8 sec/frame time-resolution. Fluorescence of GFP, mCherry and eBFP2 was excited using 488-nm, 561-nm and 405-nm lasers, respectively. Excitation power was measured and calibrated using X-Cite XR2100/XP750 Optical Power Measurement System (EXCELITAS Technologies) to keep the same experimental setting for each set of experiments. Image acquisition was started before the end of nc13 and ended after the onset of gastrulation at nc14. During imaging, data acquisition was occasionally stopped for a few seconds to correct z-position. Obtained data were concatenated and cropped into 430 x 512 pixels (*sna* shadow enhancer), 430 x 430 pixels (*Ubx* BRE) or 300 x 512 pixels (*rho* NEE) to remove nuclei outside of the expression domain. One hundred eighty timeframes staring from the entry into nc14 as defined by the progression of prior anaphase were used for subsequent image analysis. Temperature was kept in between 22.0 to 23.0 °C during imaging. For each cross, three biological replicates were taken.

### Airyscan imaging

Embryos were imaged using a Airyscan2 detector equipped in a Zeiss LSM 900 confocal microscope. Plan-Apochromat 63x / 1.4 N.A. oil immersion objective was used. Images were acquired with following settings: 944 x 944 pixels (33.8 x 33.8 μm), 16-bit depth, 41 z-slices separated by 0.2 μm. Fluorescence of GFP and mCherry was excited using 488-nm and 561-nm lasers, respectively. Excitation power was measured and calibrated using X-Cite XR2100/XP750 Optical Power Measurement System (EXCELITAS Technologies) to keep the same experimental setting throughout the experiments. Obtained images were processed using “Airyscan processing” function of Zeiss ZEN software (version 3.1) in 3D with a value of 4.6 for all the channels. Temperature was kept in between 22.0 to 23.0 °C during imaging.

### Plasmids

#### pbphi-PP7-sna shadow enhancer

A DNA fragment containing *sna* shadow enhancer was amplified using primers (5’-TTT AAG GAT CCA AGC TTG CAT TGA GGT GTT TTG-3’) and (5’-TTA AAA GAT CTG CTA GCT AAA TTC CGA TTT TTC-3’), and digested with BamHI and BglII. The resulting fragment was inserted into the unique BamHI site in pbphi-αTubulin 3 ’UTR ^21^. Subsequently, pBlueScript-24xPP7 ^3^ was digested with BamHI and BglII. The resulting fragment containing 24xPP7 was inserted into the unique BamHI site in the plasmid.

#### pbphi-No TSS-PP7-sna shadow enhancer

A DNA fragment containing a partial sequence from *lacZ* gene was amplified using primers (5’-TTT AAG CGG CCG CTC TAG AGA CGG CAG TTA TCT GGA AGA-3’) and (5’-TTA AAG GAT CCA CTT CAG CCT CCA GTA CAG C-3’), and digested with NotI and BamHI. The resulting fragment was inserted between the NotI and BamHI sites in pbphi-PP7-*sna* shadow enhancer.

#### pbphi-TSS-PP7-sna shadow enhancer

A DNA fragment containing DSCP was amplified from pbphi-DSCP-MS2-yellow ^53^ using primers (5’-TTT AAG CGG CCG CTC TAG AGA GCT CGC CCG GGG ATC GAG CGC-3’) and (5’-TTA AAG GAT CCG TTT GGT ATG CGT CTT GTG A-3’), and digested with NotI and BamHI. The resulting fragment was inserted between the NotI and BamHI sites in pbphi-PP7-*sna* shadow enhancer.

#### pbphi-TSS^weak^-PP7-sna shadow enhancer

A DNA fragment containing DSCP with Inr mutation was amplified from pbphi-DSCP_mInr_-MS2-yellow-*sna* shadow enhancer ^25^ using primers (5’-TTT AAG CGG CCG CTC TAG AGA GCT CGC CCG GGG ATC GAG CGC-3’) and (5’-TTA AAG GAT CCG TTT GGT ATG CGT CTT GTG A-3’), and digested with NotI and BamHI. The resulting fragment was inserted between the NotI and BamHI sites in pbphi-PP7-*sna* shadow enhancer.

#### pbphi-TSS^inverted^-PP7-sna shadow enhancer

A DNA fragment containing DSCP was amplified from pbphi-DSCP-MS2-yellow ^53^ using primers (5’-TTT AAG CGG CCG CTC TAG AGT TTG GTA TGC GTC TTG TGA-3’) and (5’-TTA AAG GAT CCG AGC TCG CCC GGG GAT CGA G-3’), and digested with NotI and BamHI. The resulting fragment was inserted between the NotI and BamHI sites in pbphi-PP7-*sna* shadow enhancer.

#### pbphi-DSCP-MS2-yellow-sna shadow enhancer-PP7-No TSS

pbphi-No TSS-PP7-*sna* shadow enhancer was digested with XbaI. The resulting fragment containing No TSS-PP7-*sna* shadow enhancer was inserted into the unique XbaI site in pbphi-DSCP-MS2-yellow ^53^. Orientation of the inserted DNA fragment was confirmed by sequencing.

#### pbphi-DSCP-MS2-yellow-sna shadow enhancer-PP7-TSS

pbphi-TSS-PP7-*sna* shadow enhancer was digested with XbaI. The resulting fragment containing TSS-PP7-*sna* shadow enhancer was inserted into the unique XbaI site in pbphi-DSCP-MS2-yellow ^53^ . Orientation of the inserted DNA fragment was confirmed by sequencing.

#### pbphi-DSCP-MS2-yellow-sna shadow enhancer-PP7-TSS^weak^

pbphi-TSS^weak^-PP7-*sna* shadow enhancer was digested with XbaI. The resulting fragment containing TSS^weak^-PP7-*sna* shadow enhancer was inserted into the unique XbaI site in pbphi-DSCP-MS2-yellow ^53^. Orientation of the inserted DNA fragment was confirmed by sequencing.

#### pbphi-DSCP-MS2-yellow-sna shadow enhancer-PP7-TSS^inverted^

pbphi-TSS^inverted^-PP7-*sna* shadow enhancer was digested with XbaI. The resulting fragment containing TSS^inverted^-PP7-*sna* shadow enhancer was inserted into the unique XbaI site in pbphi-DSCP-MS2-yellow ^53^. Orientation of the inserted DNA fragment was confirmed by sequencing.

#### pbphi-DSCP-MS2-yellow-TSS-PP7-sna shadow enhancer

pbphi-TSS-PP7-*sna* shadow enhancer was digested with XbaI. The resulting fragment containing TSS-PP7-*sna* shadow enhancer was inserted into the unique XbaI site in pbphi-DSCP-MS2-yellow ^53^. Orientation of the inserted DNA fragment was confirmed by sequencing.

#### pbphi-TSS-PP7-spacer

A DNA fragment containing a partial sequence from *lacZ* gene was amplified using primers (5’-GGG TTA AGC TTC TGC AGG AAT CCG ACG GGT TGT TAC T-3’) and (5’-GGG AAG CTA GCC GGA TAA ACG GAA CTG GAA A-3’), and digested with HindIII and NheI. The resulting fragment was inserted between the HindIII and NheI sites in pbphi-TSS-PP7-*sna* shadow enhancer.

#### pbphi-DSCP-MS2-yellow-sna shadow enhancer-TSS-PP7

A DNA fragment containing *sna* shadow enhancer was amplified using primers (5’-AAG GAG CTA GCG CAT TGA GGT GTT TTG TTG G-3’) and (5’-GGG GGT CTA GAT AAA TTC CGA TTT TTC TTG TCT GGG-3’), and digested with NheI and XbaI. The resulting fragment was inserted into the unique XbaI site in pbphi-DSCP-MS2-yellow ^53^. Subsequently, pbphi-TSS-PP7-spacer was digested with XbaI. The resulting fragment containing TSS-PP7-spacer was inserted into the unique XbaI site in the plasmid. Orientation of the inserted DNA fragment was confirmed by sequencing.

#### pbphi-No TSS-PP7-rho NEE

A DNA fragment containing *rho* NEE was inserted between the HindIII and NheI sites in pbphi-No TSS-PP7-*sna* shadow enhancer. Sequence of *rho* NEE is the same as one used in the previous study ^3^.

#### pbphi-TSS-PP7-rho NEE

A DNA fragment containing *rho* NEE was inserted between the HindIII and NheI sites in pbphi-TSS-PP7-*sna* shadow enhancer. Sequence of *rho* NEE is the same as one used in the previous study ^3^.

#### pbphi-DSCP-MS2-yellow-rho NEE-PP7-No TSS

pbphi-No TSS-PP7-*rho* NEE was digested with XbaI. The resulting fragment containing No TSS-PP7-*rho* NEE was inserted into the unique XbaI site in pbphi-DSCP-MS2-yellow ^53^. Orientation of the inserted DNA fragment was confirmed by sequencing.

#### pbphi-DSCP-MS2-yellow-rho NEE-PP7-TSS

pbphi-TSS-PP7-*rho* NEE was digested with XbaI. The resulting fragment containing TSS-PP7-*rho* NEE was inserted into the unique XbaI site in pbphi-DSCP-MS2-yellow ^53^. Orientation of the inserted DNA fragment was confirmed by sequencing.

#### pbphi-DSCP-MS2-yellow-rhoNEE-TSS-PP7

A DNA fragment containing *rho* NEE was amplified using primers (5’-GGG GAG CTA GCT TCC TCT GCT CAA AAT CAA A-3’) and (5’-GGA AAT CTA GAC CTC AGG TCG AGT TCC TCC A-3’), and digested with NheI and XbaI. The resulting fragment was inserted into the unique XbaI site in pbphi-DSCP-MS2-yellow ^53^. Subsequently, pbphi-TSS-PP7-spacer was digested with XbaI. The resulting fragment containing TSS-PP7-spacer was inserted into the unique XbaI site in the plasmid. Orientation of the inserted DNA fragment was confirmed by sequencing.

#### pbphi-sna shadow enhancer-PP7

A DNA fragment containing *sna* shadow enhancer was purified from pbphi-*sna* shadow enhancer ^23^ by digesting with NotI and BamHI, and inserted between the NotI and BamHI sites of pbphi-lacZ-PP7-αTubulin 3 ’UTR ^21^. Subsequently, primers (5’-GGC CGC TCT AGA CTC GAG AGT TTA-3’) and (5’-AGC TTA AAC TCT CGA GTC TAG AGC-3’) were annealed and inserted between the NotI and HindIII sites in the plasmid.

#### pbphi-DSCP-MS2-yellow-Ubx BRE-PP7

A DNA fragment containing *Ubx* BRE was amplified from genomic DNA using primers (5’-TTT TTG CTA GCA CTT CCA CTC GAA TTG CGC C-3’) and (5’-TTG GGA AGC TTT AAA TTC TCA GGC GGC ACG A-3’), and digested with NheI and HindIII. The resulting fragment was inserted between the NheI and HindIII sites in pBlueScript. Subsequently, pBlueScript-*Ubx* BRE was digested with NheI and HindIII, and the resulting DNA fragment was inserted between the NheI and HindIII site in pbphi-*sna* shadow enhancer-PP7 plasmid. Then, pbphi-*Ubx* BRE-PP7 was digested with XbaI, and the resulting DNA fragment was inserted into the unique XbaI site in pbphi-DSCP-MS2-yellow ^53^. Orientation of the inserted DNA fragment was confirmed by sequencing.

#### pbphi-DSCP-MS2-yellow-inverted Ubx BRE-PP7

A DNA fragment containing *Ubx* BRE was inserted between the NheI and HindIII sites in pbphi-No TSS-PP7-*sna* shadow enhancer. Subsequently, pbphi-inverted *Ubx* BRE-PP7 was digested with XbaI, and the resulting DNA fragment was inserted into the unique XbaI site in pbphi-DSCP-MS2-yellow ^53^. Orientation of the inserted DNA fragment was confirmed by sequencing.

#### pbphi-DSCP-MS2-yellow-inverted Ubx BRE-PP7-TSS

A DNA fragment containing *Ubx* BRE was inserted between the NheI and HindIII sites in pbphi-TSS-PP7-*sna* shadow enhancer. Subsequently, pbphi-inverted *Ubx* BRE-PP7-TSS was digested with XbaI, and the resulting DNA fragment containing inverted *Ubx* BRE-PP7-TSS was inserted into the unique XbaI site in pbphi-DSCP-MS2-yellow ^53^. Orientation of the inserted DNA fragment was confirmed by sequencing.

#### pCFD3-dU6-dl gRNA

Two DNA oligos (5’-GTC GGC AAT CAA GCG GAT AAT AA-3’) and (5’-AAA CTT ATT ATC CGC TTG ATT GC-3’) were annealed and inserted into the pCFD3-dU6:3gRNA vector (addgene# 49410) using BbsI sites.

#### pBS-dl 5’arm-GFP-3xFLAG-loxP-3xP3-dsRed-loxP-dl 3’arm

A DNA fragment containing 5’ homology arm of *dl* was amplified from genomic DNA using primers (5’-GGG GAG GTA CCG AAC CAA GAG GTG AGT TTT ATA CAC-3’) and (5’-AAA AGT CGA CCG TGG ATA TGG ACA GGT TCG ATA TCT GCA GAT CTT CCG AAT TGA GGC GCA GTA TCT GCT GAT CCT CTG AGT TTA TGT GCA CCA ACT GCC CGC TAT CGA AGC TAA GCA GAT TGC TGA GCG TTG GCG CAT TAT TAT CCG CTT GAT TGC CAG C-3’), and digested with KpnI and SalI. The resulting fragment was inserted between the KpnI and SalI sites in pBS-GFP-3xFLAG-loxP-3xP3-dsRed-loxP. Subsequently, a DNA fragment containing 3’ homology arm of *dl* was amplified from genomic DNA using primers (5’-GGG GGA CTA GTT AAT GGG CCA ACG CTC AGC AAT CTG-3’) and (5’-AAA AAG CGG CCG CGG GTG GGC AGC TTA TCC ACA-3’), and digested with SpeI and NotI. The resulting fragment was inserted between the SpeI and NotI sites in the plasmid.

#### pCFD3-dU6-zld gRNA

Two DNA oligos (5’-GTC GCA AGA GCG AGT ACG TGC AGG-3’) and (5’-AAA CCC TGC ACG TAC TCG CTC TTG-3’) were annealed and inserted into the pCFD3-dU6:3gRNA vector (addgene# 49410) using BbsI sites.

#### pBS-zld 5’arm-GFP-3xFLAG-loxP-3xP3-dsRed-loxP-zld 3’arm

A DNA fragment containing 5’ homology arm of *zld* was amplified from genomic DNA using primers (5’-TTA AAG GTA CCC GCC CTA CTC GCC CAC AGT GAG C-3’) and (5’-GGG AAG TCG ACG TAG AGC TCT ATG CTC TTC TCG ATC ATC TGA AAC TGC TCC TGC ACG TAC-3’), and digested with KpnI and SalI. The resulting fragment was inserted between the KpnI and SalI sites in pBS-GFP-3xFLAG-loxP-3xP3-dsRed-loxP. Subsequently, a DNA fragment containing 3’ homology arm of *zld* was amplified from genomic DNA using primers (5’-TTA AAT CTA GAA GGA GGA GTT TCA GAT GAT CGA G-3’) and (5’-GGG AAG CGG CCG CGC AAT GCG TTG GTC TAG TAA C-3’), and digested with XbaI and NotI. The resulting fragment was inserted between the XbaI and NotI sites in the plasmid.

### Image analysis

All the image processing methods and analysis were implemented in Fiji (https://fiji.sc), MATLAB (R2021b, MathWorks) and R (version 4.1.2).

### Segmentation of nuclei

Segmentation of nuclei was performed using Fiji. For each time point, maximum projection was obtained for all z-sections per image. His2Av-eBFP2 channel was used to segment nuclei. His2Av images were first processed with a median filter to remove salty noise and a band pass filter to remove noises below 5 pixels and above 25 pixels. Processed images were converted into binary images using an Otsús method. Touching nuclei were then segmented with a watershed algorithm. Finally, individual nuclei were separated with a Voronoi method. Resulting binary images were manually corrected to locate MS2/PP7 transcriptional dots inside the Voronoi regions.

### Tracking of nuclei

Nuclei tracking was done using custom MATLAB script, which finds the object with the minimal movement between two consecutive timeframes. First, centroids of all the segmented nuclei were determined as a location of each nucleus for all the timeframes. For each timeframe, Euclidean distances between centroids in the current and the previous timeframe were determined. Nucleus with the minimum Euclidean distance was considered as the same lineage. When a nucleus touched the edge of the image or moved larger than the length of nucleus, corresponding lineage was excluded from the analysis.

### Recording of MS2 and PP7 signal

Raw images were maximum projected for each timeframe and used to record MS2 and PP7 fluorescence intensities. Using segmented regions, fluorescence intensities within each nucleus were extracted. Signals of MS2 and PP7 transcription dots were determined by taking integral of a 5 x 5 pixels region centering the brightest pixel within a nucleus after subtracting median fluorescence intensity within a nucleus as a background. Subsequently, minimum intensities were determined for individual trajectories and subtracted to make the baseline zero.

### Detection of transcriptional bursting

First, each trajectory was smoothed by lowess function within a window of 10 timeframes in R. When a nucleus had above-threshold transcription activity, burst was considered to be started. Burst was considered to be ended when the intensity dropped below the threshold. When two connective bursts are touching with each other, they were divided at the timeframe where signal intensity successively decreased from prior 2 timeframes and increased in subsequent 2 timeframes. When burst duration is less than 5 timeframes, it was considered as a false-positive. It was confirmed that this filtering method does not affect quantification of burst property ^25^. From each nucleus, total number of bursts was determined as the burst frequency. Cumulative fraction of active nuclei (Figure S2) was determined by calculating the faction of nuclei that experienced burst production after the entry into nc14. In the analysis of *rho* NEE reporters (Figure S3), nuclei located at the middle of expression domain were used.

### Quantification of burst size and total output

Total output was determined for each nucleus by integrating signal intensities at all the time points using raw trajectory. False-colored images were generated using segmentation mask at the 50th timeframe of the data (∼14 minutes after the entry into nc14). They were colored with the pixel intensity proportional to the total output in corresponding nuclei. Burst size was determined by integrating signal intensities during individual bursts using raw trajectory.

### Radial analysis of Airyscan images

Segmentation of nuclei was performed as described above. GFP channel was used for segmentation. Using a band pass filter, binarized objects with the size between 50 to 200 pixels were obtained. Resulting segmentation images were then used to compute a 3D voxel segment region for each nucleus. To quantify local GFP intensity centering the PP7 transcription site, XYZ coordinate of the brightest PP7 pixel intensity was determined in each 3D-segmented nucleus. Dl-GFP signal at 29 x 29 pixels (∼1.0 x 1.0 μm) or Zld-GFP signal at 41 x 41 pixels (∼1.4 x 1.4 μm) region centering the brightest PP7 signal was measured on the same z-plane. As a control, Dl-GFP or Zld-GFP signal centering the random XY coordinate within the segmented region on the same z-plane was measured. To analyze nuclei that reliably contain active PP7 transcription site, nuclei with top 15% of integrated PP7 signal were pooled from all the analyzed embryos. Then, obtained images were separated into bins based on distance from the center in a pixel increment (1 pixel; 35.5 nm). To determine relative enrichment of transcription factors (Figure 5C, Figure S4A), Dl-GFP or Zld-GFP intensities were then divided by mean intensity of the circumscribed or inscribed bin area, respectively. For quantification of radial profile of Dl-GFP and Zld-GFP distribution (Figure 5D, Figure S4B), mean intensity of each bin was divided by the mean of endmost bin and then standard error of the mean was calculated for each bin.

### Quantification of TSS score on developmental enhancers

Publicly available 5’ CAGE-seq data of 2-4h WT *Drosophila melanogaster* embryos (E-MTAB-4787) ^58^ were used to quantify the level of non-coding transcription at the enhancer regions. Sequenced reads were mapped to the BDGP *Drosophila melanogaster* genome release 3 (dm3) using HISAT2 (version 2.2.0) ^59^. Samtools (version 1.10) and bedtools (version 2.29.2) were used to convert files to bed format. To exclude TSS signals derived from the coding regions, reads aligned within a window of 50 bp from any exonic sequences were removed using “bedtools intersect” function. Remaining reads were integrated into a single file. Next, to quantify TSS score on enhancers, “bedtools coverage-counts” function was used to calculate the number of reads on developmental enhancers that were reliably assigned to their target genes in previous study ^60^. For quantitative comparison of TSS score, the number of reads was normalized to the size of each enhancer.

