## Supplementary figures and images for "Dynamic interplay between non-coding enhancer transcription and gene activity in development"

### Supplemental Figure

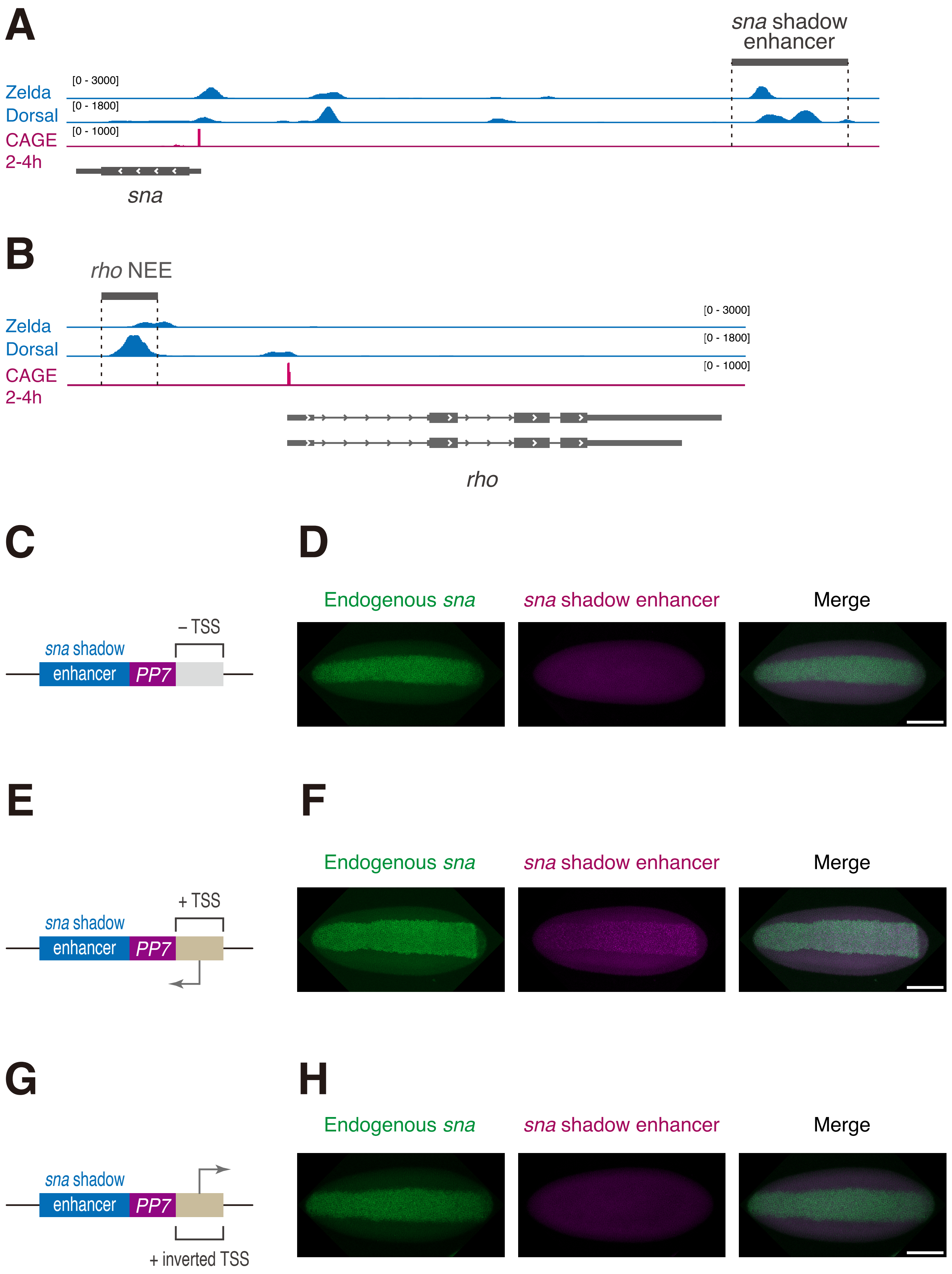

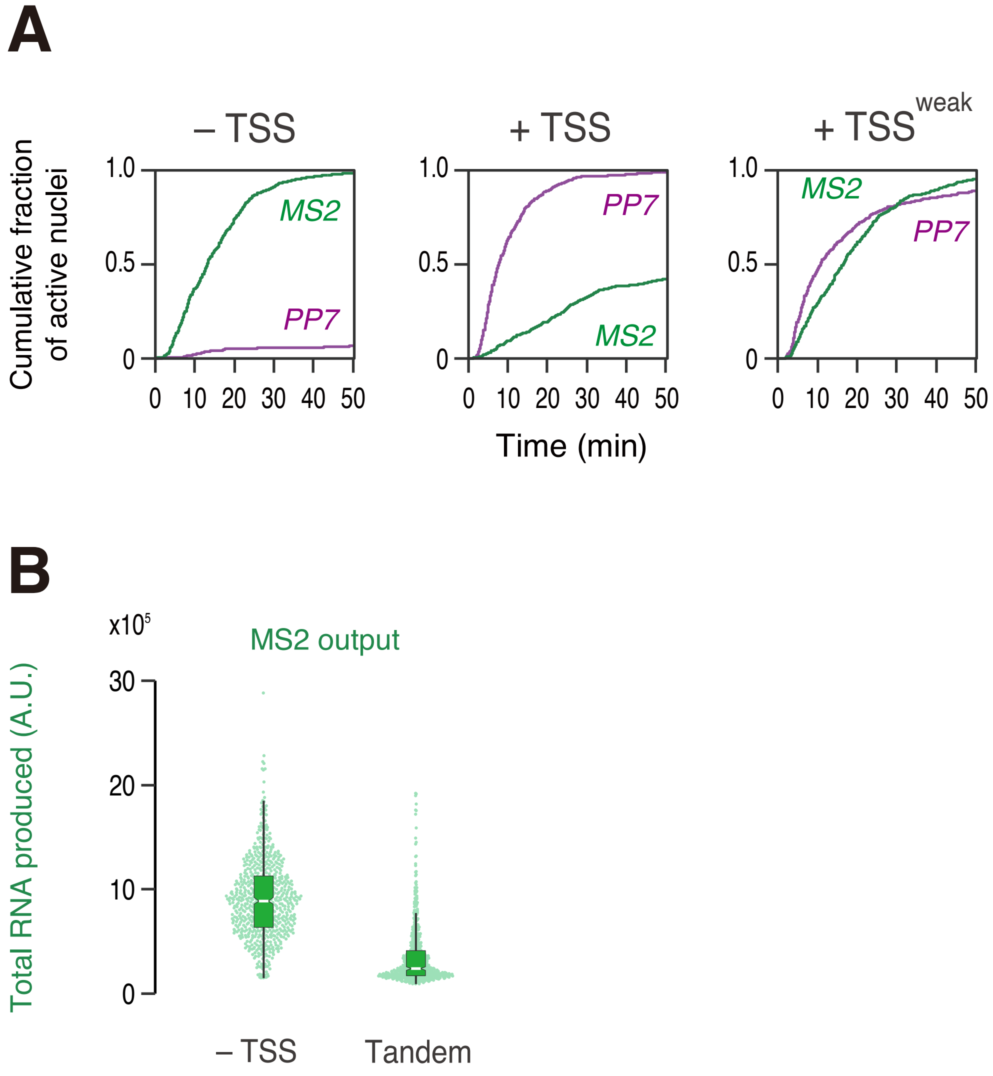

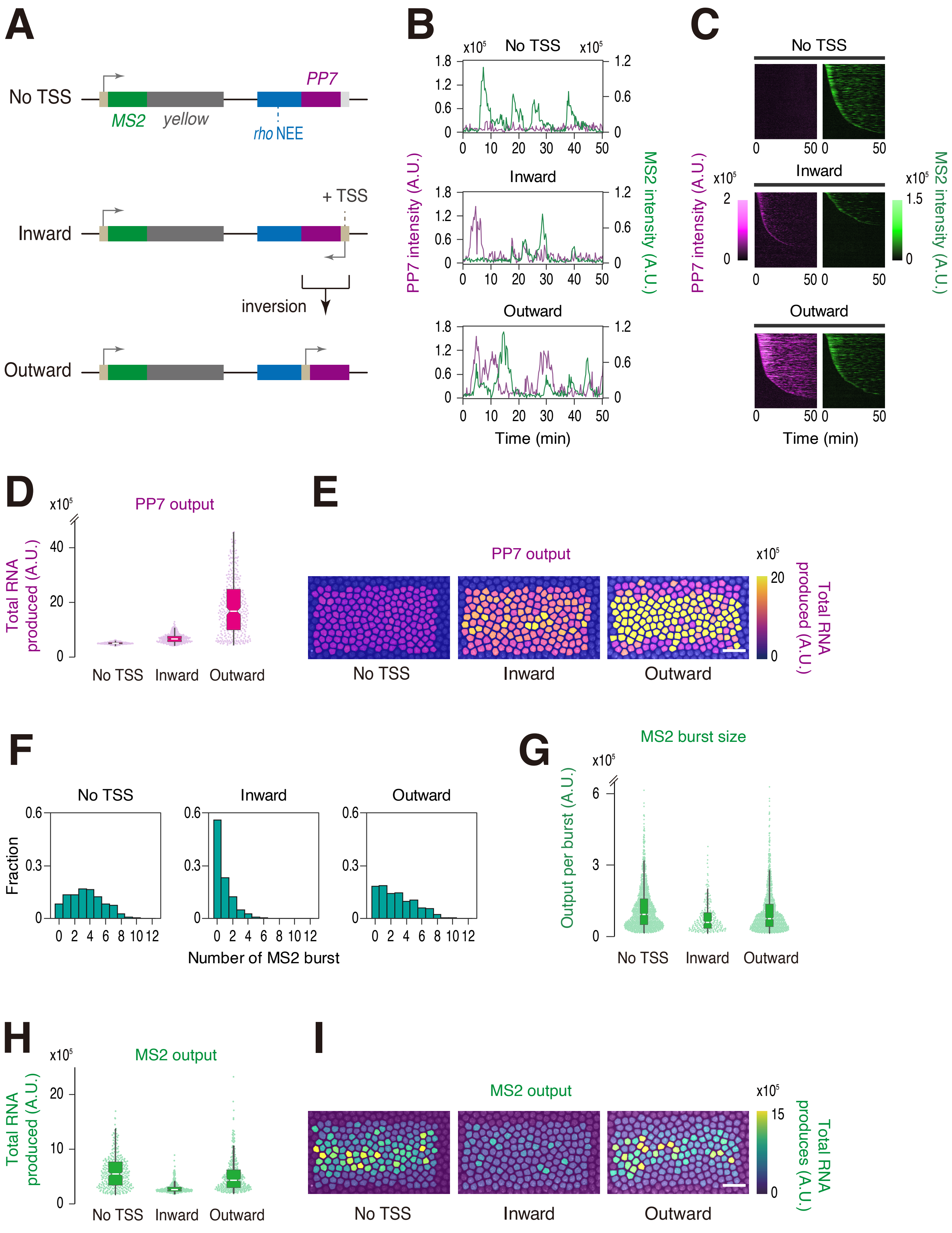

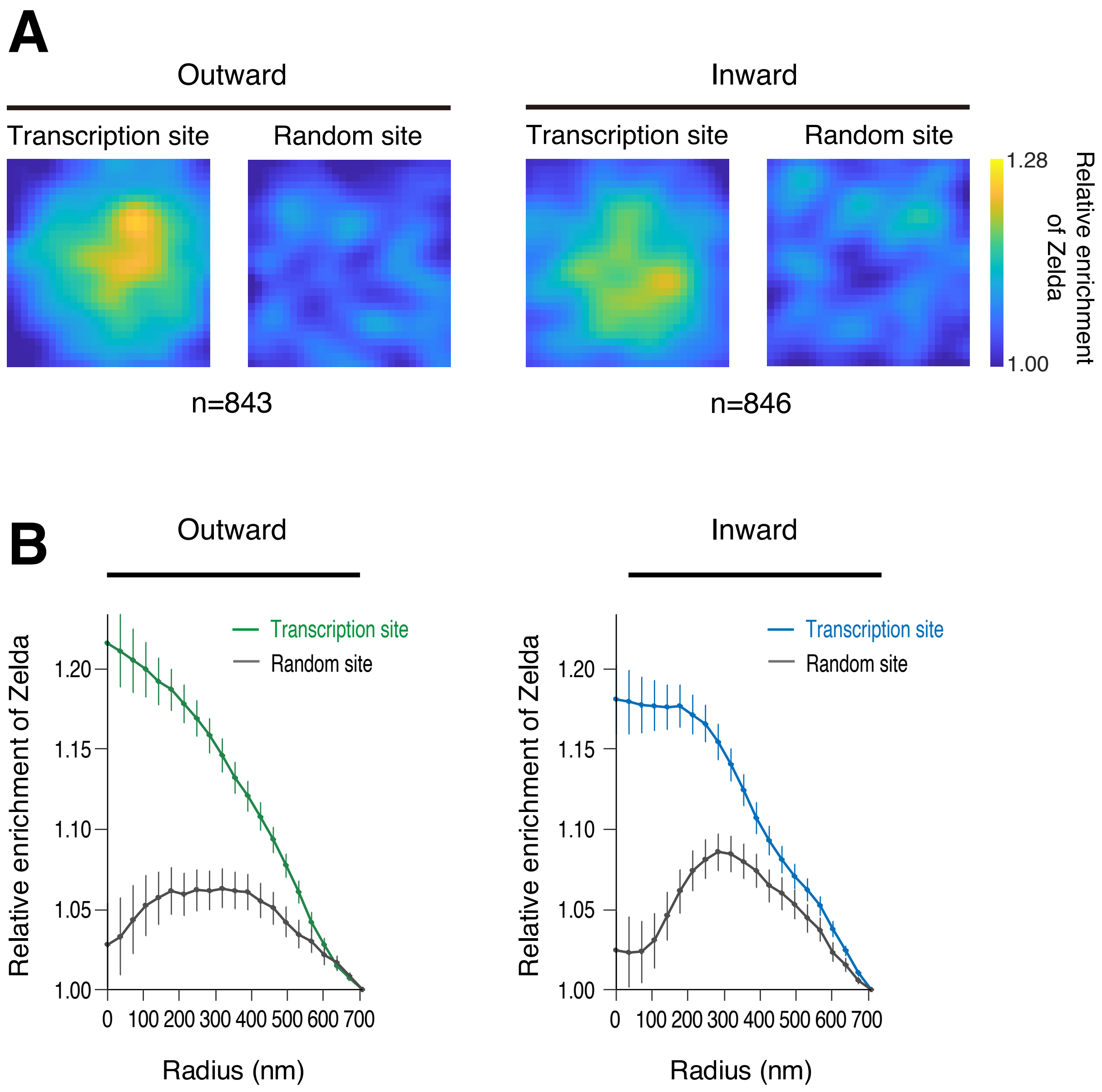

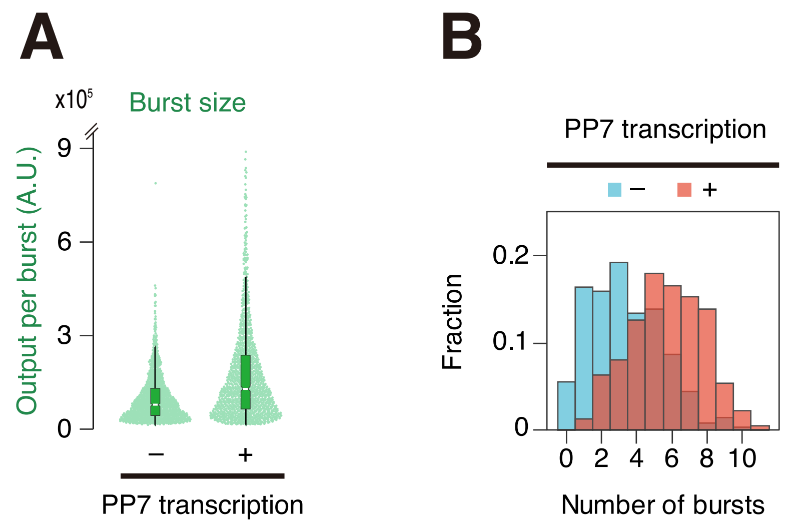

Hamamoto *et al.*, Figure S6

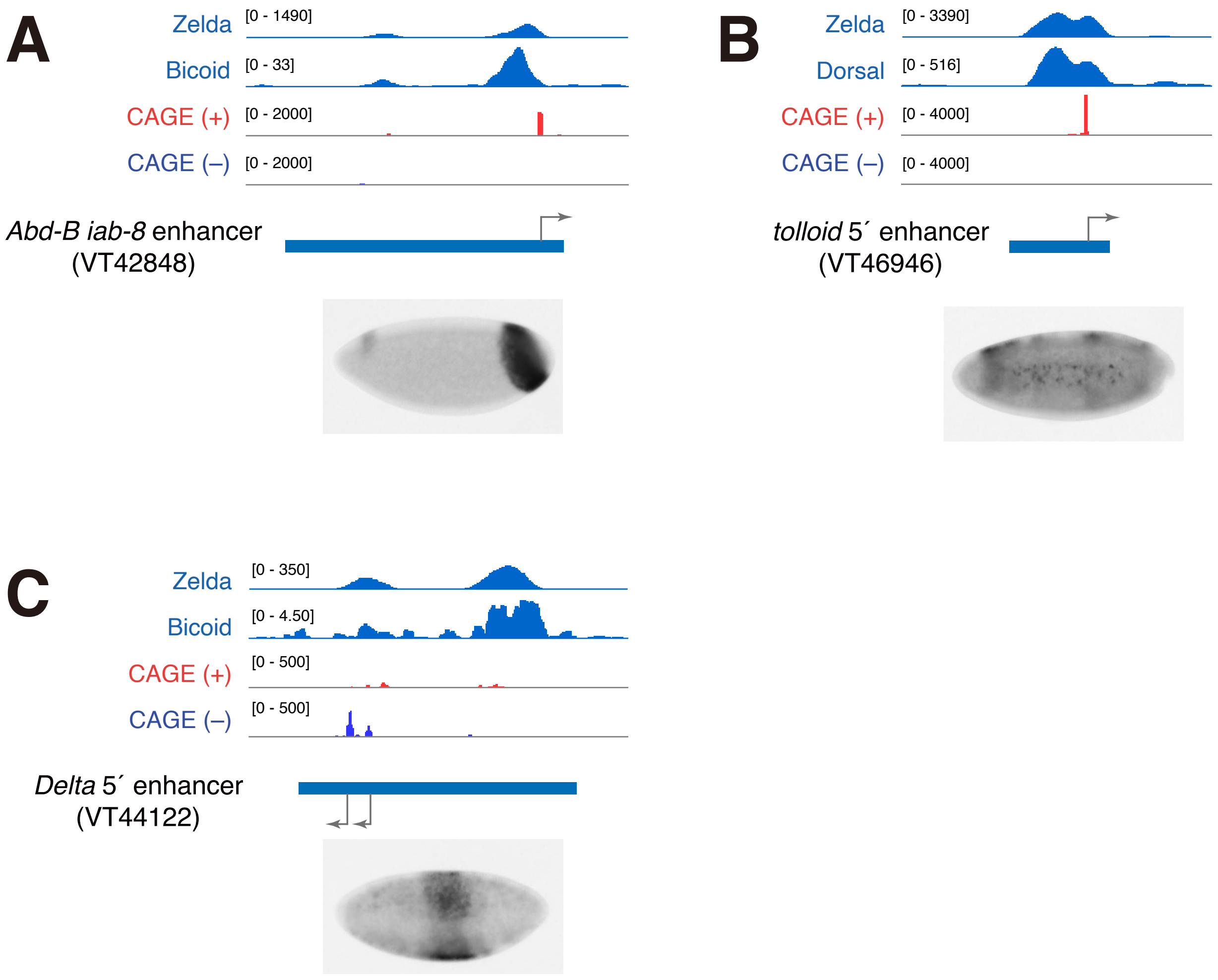
