## Supplemental Figure Legends for "Dynamic interplay between non-coding enhancer transcription and gene activity in development"

**Figure S1. Induction of unidirectional non-coding enhancer transcription at the synthetic locus. Related to Figure 1.**

(A) Organization of the endogenous *sna* locus. Zelda ChIP-seq data from nc13 WT embryos (GSM763061) [1], Dorsal ChIP-seq from 2- to 4-h WT embryos (GSM1341814) [2], CAGE-seq data from 2- to 4-h WT embryos (ERR1425056) [3] were visualized with Integrative Genomics Viewer.

(B) Organization of the endogenous *rho* locus. Zelda ChIP-seq data from nc13 WT embryos (GSM763061) [1], Dorsal ChIP-seq from 2- to 4-h WT embryos (GSM1341814) [2], CAGE-seq data from 2- to 4-h WT embryos (ERR1425056) [3] were visualized with Integrative Genomics Viewer.

(C) Schematic representation of the PP7-enhancer cassette without TSS.

(D) Fluorescent *in situ* hybridization using probes against endogenous *sna* gene (left) and *sna* shadow enhancer (middle). nc14 embryos containing the PP7-enhancer cassette without TSS were analyzed. The image is oriented with anterior to the left and ventral view facing up. Scale bar indicates 100 μm.

(E) Minimal core promoter motifs were placed adjacent to the enhancer to drive non-coding enhancer transcription.

(F) Fluorescent *in situ* hybridization using probes against endogenous *sna* gene (left) and *sna* shadow enhancer (middle). nc14 embryos containing the PP7-enhancer cassette fused with promoter motifs were analyzed. The image is oriented with anterior to the left and ventral view facing up. Scale bar indicates 100 μm.

(G) Inverted promoter motifs were fused with the enhancer.

(H) Fluorescent *in situ* hybridization of embryos using probes against endogenous *sna* gene (left) and *sna* shadow enhancer (middle). nc14 embryos containing the PP7-

enhancer cassette fused with inverted promoter motifs were analyzed. The image is oriented with anterior to the left and ventral view facing up. Scale bar indicates 100 μm.

**Figure S2. Activation kinetics of enhancer and target gene transcription. Related to Figure 1 and 3.**

(A) Cumulative fraction of actively transcribing nuclei. A total of 676, 719, and 700 ventral-most nuclei, respectively, were analyzed from three independent embryos for the reporter locus containing – TSS (top), + TSS (middle), or + $TSS^{weak}$ (bottom) at the enhancer region.

(B) Boxplot showing the distribution of total output of MS2 transcription. A total of 676 and 638 ventral-most nuclei, respectively, were analyzed from three independent embryos for the reporter locus without intergenic TSS or driving non-coding enhancer transcription in a tandem orientation. The box indicates the lower (25%) and upper (75%) quantile and the white line indicates the median. Whiskers extend to the most extreme, non-outlier data points. Plot of – TSS is the same as the plot shown in Figure 1H. Plot of Tandem is the same as the plot shown in Figure 3G.

**Figure S3. Non-coding transcription of *rho* NEE attenuates target gene transcription. Related to Figure 1 and 4.**

(A) The *MS2-yellow* reporter gene was placed under the control of the *rho* NEE fused with 24x PP7 repeats (top). Minimal core promoter motifs were placed adjacent to the enhancer to drive non-coding transcription in an inward orientation (middle). PP7 transcription unit was inverted to drive non-coding transcription in an outward orientation (bottom).

(B) Representative trajectories of transcriptional activities of the reporter locus with No TSS (top), inward PP7 transcription (middle), or outward PP7 transcription (bottom).

A.U.; arbitrary unit.

(C) MS2 and PP7 trajectories for all analyzed nuclei. Each row represents the MS2 or PP7 trajectory for a single nucleus. A total of 331, 313, and 345 nuclei, respectively, were analyzed from three independent embryos for the reporter locus with No TSS (top), inward PP7 transcription (middle), or outward PP7 transcription (bottom). Nuclei were ordered by the onset of MS2 or PP7 transcription in nc14, separately. The same number of nuclei were analyzed hereafter.

(D) Boxplot showing the distribution of total output of PP7 transcription. The box indicates the lower (25%) and upper (75%) quantile and the white line indicates the median. Whiskers extend to the most extreme, non-outlier data points. The double hash mark on the y-axis indicates that > 99% of the data points are presented.

(E) Each nucleus was colored with respect to the total output of PP7 transcription in the representative embryos. The maximum projected image of His2Av-eBFP2 is shown in gray. The image is oriented with anterior to the left. Scale bar indicates 20 μm.

(F) Histograms showing the distribution of MS2 burst frequency.

(G) Boxplot showing the distribution of MS2 burst size. The box indicates the lower (25%) and upper (75%) quantile and the white line indicates the median. Whiskers extend to the most extreme, non-outlier data points. A total of 1159, 243, and 964 MS2 bursts, respectively, were analyzed for the reporter locus with No TSS, inward PP7 transcription, or outward PP7 transcription. The double hash mark on the y-axis indicates that >99% of the data points are presented.

(H) Boxplot showing the distribution of total output of MS2 transcription. The box indicates the lower (25%) and upper (75%) quantile and the white line indicates the median. Whiskers extend to the most extreme, non-outlier data points.

(I) Each nucleus was colored with respect to the total output of MS2 transcription in the representative embryos. The maximum projected image of His2Av-eBFP2 is shown in gray. The image is oriented with anterior to the left. Scale bar indicates 20 μm.

**Figure S4. Enhancer self-transcription decreases local concentration of pioneering transcription factor Zld. Related to Figure 5.**

(A) Heatmaps showing the averaged distribution of Zelda-GFP centering the PP7 transcription site or random site. A total of 843 and 846 PP7-transcribing nuclei, respectively, were obtained from 50 independent embryos for the reporter locus driving PP7 transcription in an outward (left) or an inward orientation (right).

(B) Radial profiles of the averaged Zelda-GFP distribution shown in (A). Error bar represents standard error of the mean.

**Figure S5. Non-coding BRE transcription correlates with the efficiency of burst induction. Related to Figure 6.**

(A) Boxplot showing the distribution of MS2 burst size. The box indicates the lower (25%) and upper (75%) quantile and the white line indicates the median. Whiskers extend to the most extreme, non-outlier data points. A total of 1195 and 1278 MS2 bursts, respectively, were analyzed from nuclei grouped by the absence or presence of PP7 transcription from unmodified BRE (Figure 6C; top). The double hash mark on the y-axis indicates that >99% of the data points are presented.

(B) Histograms showing the distribution of MS2 burst frequency. A total of 359 and 223 nuclei grouped by the absence or presence of PP7 transcription from unmodified BRE (Figure 6C; top), were analyzed respectively.

**Figure S6. Organization of other transcribing enhancers identified in this study. Related to Figure 6.**

(A) Organization of the endogenous *Abd-B iab-8* enhancer. Zelda ChIP-seq data from nc13 WT embryos (GSM763061) [1], Bicoid-GFP ChIP-seq from nc14 WT embryos (GSE86966) [4], processed 2- to 4-h WT CAGE-seq data (E-MTAB-478) [3] after excluding reads mapped to the coding regions were visualized with Integrative Genomics Viewer. A picture showing the activity of *Abd-B iab-8* enhancer (VT42848) was taken from Fly Enhancers [5].

(B) Organization of the endogenous *tolloid* 5′ enhancer. Zelda ChIP-seq data from nc13 WT embryos (GSM763061) [1], Dorsal ChIP-seq from 2- to 4-h WT embryos (GSM1341814) [2], processed 2- to 4-h WT CAGE-seq data (E-MTAB-478) [3] after excluding reads mapped to the coding regions were visualized with Integrative Genomics Viewer. A picture showing the activity of *tolloid* 5′ enhancer (VT46946) was taken from Fly Enhancers [5].

(C) Organization of the endogenous *Delta* 5′ enhancer. Zelda ChIP-seq data from nc13 WT embryos (GSM763061) [1], Bicoid-GFP ChIP-seq from nc14 WT embryos (GSE86966) [4], processed 2- to 4-h WT CAGE-seq data (E-MTAB-478) [3] after excluding reads mapped to the coding regions were visualized with Integrative Genomics Viewer. A picture showing the activity of *Delta* 5′ enhancer (VT44122) was taken from Fly Enhancers [5].

### Supplemental Movie Legends

**Movie S1. Live-imaging of embryos driving non-coding enhancer transcription *in cis*.**

MS2/PP7 two-color live-imaging of embryos with the reporter locus containing – TSS (top), + TSS (middle), or $TSS^{weak}$ (bottom) at the enhancer region. The maximum projected images of PP7, MS2 and His2Av signal are shown in magenta, green and grey, respectively. The image is oriented with anterior to the left and ventral view facing up. Scale bar indicates 30 μm.

**Movie S2. Live-imaging of embryos driving non-coding enhancer transcription *in trans*.**

MS2/PP7 two-color live-imaging of embryos lacking (top) or containing (bottom) the PP7 allele. The maximum projected images of PP7, MS2 and His2Av signal are shown in magenta, green and grey, respectively. The image is oriented with anterior to the left and ventral view facing up. Scale bar indicates 30 μm.

**Movie S3. Live-imaging of embryos driving non-coding enhancer transcription in a tandem orientation.**

MS2/PP7 two-color live-imaging of embryos with the reporter locus driving non-coding enhancer transcription in a convergent (top) or a tandem orientation (bottom). The maximum projected images of PP7, MS2 and His2Av signal are shown in magenta, green and grey, respectively. The image is oriented with anterior to the left and ventral view facing up. Scale bar indicates 30 μm. Movie of Convergent is the same as the movie of + TSS shown in Movie S1.

**Movie S4. Live-imaging of embryos driving PP7 transcription in an outward orientation.**

MS2/PP7 two-color live-imaging of embryos driving PP7 transcription in an inward (top) or an outward orientation (bottom). The maximum projected images of PP7, MS2 and His2Av signal are shown in magenta, green and grey, respectively. The image is oriented with anterior to the left and ventral view facing up. Scale bar indicates 30 μm.

**Movie S5. Calculation of average Dorsal distribution around non-inhibitory PP7 transcription site.**

Top panel shows images of the PP7 signal, corresponding Dorsal-GFP signal in the same window, and Dorsal-GFP signal at a random site in the same nucleus. Middle panel shows a running average of the images in the top panel. Bottom panel shows the corresponding running average radial profile. A total of 839 nuclei from 50 independent embryos were analyzed.

**Movie S6. Calculation of average Dorsal distribution around inhibitory PP7 transcription site.**

Top panel shows images of the PP7 signal, corresponding Dorsal-GFP signal in the same window, and Dorsal-GFP signal at a random site in the same nucleus. Middle panel shows a running average of the images in the top panel. Bottom panel shows the corresponding running average radial profile. A total of 852 nuclei from 50 independent embryos were analyzed.

**Movie S7. Live-imaging of embryos driving non-coding enhancer transcription from *Ubx* BRE.**

MS2/PP7 two-color live-imaging of embryos with the reporter locus driving PP7 transcription from BRE. The maximum projected images of PP7, MS2 and His2Av signal are shown in magenta, green and grey, respectively. The image is oriented with anterior to the left and lateral view facing up. Scale bar indicates 30 μm.
