## Supplemental Table 1 for "Dynamic interplay between non-coding enhancer transcription and gene activity in development"

| Location of enhancer | FlyEnhancer ID | Target gene | TSS score (reads/kb) | Enhancer size (kb) | Cumulative fraction of TSS score |
|---|---|---|---|---|---|
| chr3L:9,797,330-9,797,598 | VT28267 | *Ilp4* | 6007.5 | 0.268 | 100 |
| chr3R:20,574,703-20,575,504 | VT46946 | *tld* | 5048.7 | 0.801 | 91.55642245 |
| chr3R:20,997,630-20,999,927 | VT47167 | *danr* | 3410.5 | 2.297 | 84.46041528 |
| chr3R:12,524,782-12,526,937 | VT42746 | *Ubx* | 3116.0 | 2.155 | 79.66685728 |
| chr3R:12,747,382-12,749,489 | VT42849 | *Abd-B* | 2499.3 | 2.107 | 75.28726021 |
| chr3R:17,201,451-17,203,140 | VT45189 | *mod(mdg4)* | 2016.0 | 1.689 | 71.77447388 |
| chr2L:10,388,040-10,390,080 | VT5287 | *da* | 1903.4 | 2.04 | 68.94097607 |
| chr3L:14,749,272-14,751,412 | VT30868 | *Trl* | 1390.2 | 2.14 | 66.26567516 |
| chr3R:21,008,302-21,010,513 | VT47173 | *danr* | 1165.1 | 2.211 | 64.31174691 |
| chr3R:12,526,519-12,528,610 | VT42747 | *Ubx* | 1122.0 | 2.091 | 62.6742046 |
| chr3R:12,075,311-12,077,425 | VT42491 | *tara* | 1034.5 | 2.114 | 61.09728558 |
| chr3L:20,396,551-20,398,733 | VT33785 | *trbl* | 999.1 | 2.182 | 59.64323601 |
| chr2R:11,016,930-11,019,079 | VT17013 | *chn* | 910.7 | 2.149 | 58.23900951 |
| chr2L:2,752,765-2,754,884 | VT1404 | *Bacc* | 870.7 | 2.119 | 56.95906889 |
| chr2L:11,794,041-11,794,478 | VT6030 | *crol* | 691.1 | 0.437 | 55.735296 |
| chr2R:19,467,816-19,469,873 | VT21426 | *apt* | 664.1 | 2.057 | 54.76397915 |
| chr2R:2,583,575-2,585,628 | VT12642 | *Rab2* | 638.1 | 2.053 | 53.83061348 |
| chr3L:20,398,155-20,400,412 | VT33786 | *trbl* | 637.1 | 2.257 | 52.93376772 |
| chr3R:12,073,519-12,075,931 | VT42490 | *tara* | 610.3 | 2.412 | 52.03827359 |
| chr3R:12,745,522-12,747,731 | VT42848 | *Abd-B* | 608.4 | 2.209 | 51.18051333 |
| chr3R:18,352,267-18,354,359 | VT45801 | *CG5346* | 600.4 | 2.092 | 50.3253699 |
| chr3L:8,645,778-8,647,917 | VT27671 | *h* | 596.5 | 2.139 | 49.48152355 |
| chr3L:20,688,420-20,690,975 | VT33934 | *kni* | 576.1 | 2.555 | 48.64307715 |
| chr2L:2,162,032-2,164,151 | VT1082 | *aop* | 554.0 | 2.119 | 47.83332461 |
| chr2R:20,400,628-20,402,887 | VT21906 | *Letm1* | 504.6 | 2.259 | 47.05462034 |
| chr2L:2,163,674-2,165,773 | VT1083 | *aop* | 455.5 | 2.099 | 46.34533002 |
| chr3R:7,604,792-7,606,922 | VT40165 | *Lk6* | 452.1 | 2.13 | 45.70518132 |
| chr3R:2,684,413-2,686,617 | VT37567 | *ftz* | 444.6 | 2.204 | 45.06973027 |
| chr2R:10,060,721-10,062,851 | VT16530 | *cg* | 416.9 | 2.13 | 44.44477361 |
| chr2L:8,844,948-8,847,235 | VT4450 | *SoxN* | 405.3 | 2.287 | 43.85881252 |
| chr3R:15,166,068-15,168,266 | VT44122 | *Dl* | 397.2 | 2.198 | 43.2891089 |
| chr2R:1,597,188-1,599,358 | VT12230 | *ap* | 392.6 | 2.17 | 42.73086759 |
| chr2L:21,843,469-21,845,803 | VT11145 | *tsh* | 384.7 | 2.334 | 42.17902492 |
| chr3R:15,140,147-15,142,388 | VT44107 | *Dl* | 384.2 | 2.241 | 41.63825703 |
| chr2L:21,841,894-21,844,018 | VT11144 | *tsh* | 374.8 | 2.124 | 41.09825336 |
| chr3L:9,013,475-9,016,012 | VT27864 | *Doc1* | 368.9 | 2.537 | 40.57151619 |
| chrX:5,479,796-5,481,897 | VT57351 | *Vsx1* | 368.4 | 2.101 | 40.052966 |
